# Coupling comprehensive pesticide-wide association study to iPSC dopaminergic screening identifies and classifies Parkinson-relevant pesticides

**DOI:** 10.1101/2022.02.06.479305

**Authors:** Kimberly C Paul, Richard C Krolewski, Edinson Lucumi Moreno, Jack Blank, Kris Holton, Tim Ahfeldt, Melissa Furlong, Yu Yu, Myles Cockburn, Laura K Thompson, Jeff Bronstein, Lee L. Rubin, Vikram Khurana, Beate Ritz

**Author notes:** co-first authors (Kimberly C Paul, Richard C Krolewski). co-corresponding PIs.

## Abstract

Parkinson’s disease (PD) is a complex, multi-factorial neurodegenerative disease, known to involve genetic, aging-related components, but also to be highly sensitive to environmental factors. In particular, ample evidence links pesticides to PD etiology. Here, establishing a field-to-bench paradigm, we have combined record-based exposure assessment in a population-based epidemiologic study of PD with testing in dopaminergic neurons produced from iPSCs to further identify and classify PD-relevant pesticides. First, agricultural pesticide-application records in California enabled us to investigate exposure to nearly 300 specific pesticides and PD risk in a comprehensive, pesticide-wide association study (PWAS). We implicated long-term exposure to 53 pesticide active ingredients in PD risk and identified their relevant co-exposure profiles. Second, to identify which of these pesticides might contribute to PD through direct effects on dopaminergic neurons, we employed a live-cell imaging screening paradigm in which neurons, definitively identified with a tyrosine hydroxylase reporter, were exposed to 43 of the high-risk pesticides. Using detailed morphometric measures, we found 10 pesticides were directly toxic to these neurons. Further, we analyzed pesticides typically used in combinations in cotton farming. Among these “cotton cluster” pesticides, co-exposures resulted in markedly greater toxicity than any single pesticide. Trifluralin was a pivotal driver of toxicity to dopaminergic neurons and led to marked mitochondrial dysfunction. Our field-to-bench paradigm may prove useful to mechanistically dissect pesticide exposure implicated in PD risk, and guide agricultural policy in the future.

## INTRODUCTION

Parkinson’s disease (PD) is a complex, multi-factorial neurodegenerative disease, known to involve genetic, environmental, and aging-related components [1]. The hallmark pathology of PD is aggregation of the protein alpha-synuclein (α-syn) in Lewy bodies in specific midbrain dopaminergic (mDA) neurons. Considerable advances have been made in identifying environmental risk factors for PD, including ample evidence linking pesticides in general to PD etiology[2]. Yet, aside from the widely accepted model organism and *in vitro* experimental use of rotenone, studies have been limited in addressing the specific effects of most common, widely-used pesticides and few address co-exposures[3]. Fewer studies still have delved into whether pesticides may directly exert effects on dopaminergic neurons[4]. In the state of California, which is the largest agricultural producer and exporter in the United States, there are currently 13,754 pesticide products with 1,066 different active ingredients registered for use[5]. While pesticides are important components of modern commercial agriculture that help maximize food production, the majority of pesticides that are applied at industrial scales have not been adequately assessed for their potential role in PD, let alone their mechanisms of action. Consequently, a wider screen of commercially applied pesticides in relation to PD, coupled to analysis in tractable disease-relevant models, may identify new targets, provide mechanistic insights, and help revise priorities for research and public health policy.

We therefore developed a field-to-bench paradigm, coupling systematic epidemiologic screening with direct testing in neurons, to assess and mechanistically dissect pesticide-PD relationships. First, we performed a pesticide-wide association study (PWAS). Specifically, we established a record-based exposure assessment approach using agricultural pesticide application records in California to comprehensively investigate long-term ambient pesticide exposure in relation to PD risk. This allowed us to screen nearly 300 specific pesticide active ingredients in an untargeted manner, without relying on self-reported exposure or pre-selection of specific pesticides and to assess their association with PD in a population-based study. By analyzing all pesticides with an adequate exposure prevalence across the study period in a hypothesis-free manner (n=288 pesticides, 1974-2015), we sought to agnostically implicate pesticides with PD. Then, we followed this with a systematic analysis of the pesticide hits in dopaminergic neurons derived from PD patient induced pluripotent stem cells (iPSC). This cellular platform enabled us to directly test whether pesticides identified via our PWAS exert an adverse effect on PD-patient derived mDA neurons.

We chose mDA neurons produced from induced pluripotent stem cells (iPSCs) as our model-system because they represent an unprecedented tool for personalized *in vitro* disease modeling, including for central nervous system (CNS) cells[6–9]. Moreover, human iPSC model systems express proteins at endogenous levels and harbor disease-relevant pathologies. For instance, PD iPSC-derived neurons show defective mitochondrial function, ER-to-Golgi trafficking, reduced protein synthesis, increased nitrosative stress, and deficient survival over time, and disease-associated changes observed *in vitro* can be traced back to individual patients. Improved protocols for differentiation of iPSC into midbrain dopaminergic neurons exist[7, 8, 10–14]. But heterogeneity of differentiation line-to-line, clone-to-clone and experiment-to-experiment remains a challenge. To create a tractable tool useful for large scale screening purposes, we have developed a bright red fluorescent reporter to mark neurons expressing tyrosine hydroxylase, the rate-limiting enzyme in dopamine synthesis[15]. This system enabled us to study the direct effects of pesticides on mDA neurons in isolation, free from the presence of other cell types.

Commercial pesticide use remains a cornerstone of modern agriculture. Thus, population-based human research that comprehensively addresses pesticide exposures in agricultural regions as it relates to major human diseases like PD is urgently needed. Equally urgent is laboratory-based analysis that allows us to understand the mechanistic basis of these associations. The epidemiologic and iPSC-based approaches interwoven in the field-to-bench paradigm we outline here enable us to do just that. Among the multitude of potentially PD-relevant pesticides we identified, we were able to pinpoint ten that were directly toxic to mDA neurons. Further, co-exposure data common in agricultural practices allowed us to develop co-exposure paradigms “in the dish” to test whether combinations of pesticides lead to greater, synergistic toxicity. For example, from pesticides used in combination in cotton agriculture, we identified trifluralin together with other commonly co-applied pesticides as being significantly more toxic to mDA neurons than any of the cotton applied pesticides alone. We further attributed the trifluralin-driven neurotoxicity to mitochondrial dysfunction in those neurons. In time, our approach will enable us to further track such cellular pathologies back to epidemiologic and environmental data, to mechanistically understand the individual and combined effects of pesticides, and to hopefully inform the judicious use of pesticides in agriculture.

## RESULTS

### Population-Based Study Overview

Since 1974, California law mandates the recording of commercial pesticide use to the pesticide use report (PUR) database, documenting nearly 50 years of agricultural application of hundreds of pesticide active ingredients across the state. This database records the location of pesticide applications, which can be linked to the Public Land Survey System (PLSS), the poundage, type of crop, and acreage subjected to pesticide application, as well as the method and date of application. We have designed a geospatial algorithm which combines this database with maps of land-use and crop cover, providing a digital representation of historic land-use, to determine pesticide applications at specific agricultural sites each year since 1974[16]. This geographic information systems (GIS)-based algorithm allowed us to determine for each individual pesticide active ingredient in the PUR, the reported pounds of each pesticide applied per acre within a 500m buffer around specific locations, such as addresses, yearly since 1974. We applied this system to lifetime residential and workplace address histories from participants of the Parkinson’s Environment and Genes (PEG) study (n=829 PD patients; n=824 controls).

PEG is a population-based PD case-control study conducted in three agricultural counties in Central California (Kern, Fresno, and Tulare)[17]. Participants were recruited and enrolled in two waves: wave 1 (PEG1): 2000-2007, n=357 PD patients, n=400 population-based controls; wave 2 (PEG2): 2009-2015, n=472 PD patients, n=424 population-based controls. Patients were first identified from neurology offices, hospital records, and public service announcements (PEG1) and then through the California PD registry pilot program (PEG2). Population controls were randomly sampled from Medicare records first and later from county tax assessor records and approached by mail, phone, and in-person. Patients were enrolled early in disease course (mean PD duration at baseline 3.0 years (SD=2.6)), and all were seen by UCLA movement disorder specialists (lead by J.B.) for *in-person* neurologic exams and confirmed as having clinically-defined, idiopathic PD [18]. The two distinct waves of PEG (PEG1 and PEG2) allowed us to use the first wave as the discovery data set and the second to replicate the findings.

For each pesticide active ingredient, hereafter referred to as “pesticide”, in the PUR and each PEG participant, we determined the average pounds of pesticide applied per acre per year within a 500m buffer of each residential and workplace address over the study window (1974 to 10 years prior to index date, which was PD diagnosis for patients or interview date for controls) We weighed the total poundage by the proportion of acreage treated (lbs/acre). This approach created one summary estimate of the average pounds of pesticide applied per acre per year within the 500m buffer for each pesticide.

### Extent of agricultural pesticide applications in study area

From the start of pesticide use report (PUR) tracking in 1974 to 2017, there have been approximately 5.9 million PUR records in the tri-county study area (Kern, Fresno, and Tulare), documenting the application of 1,355 unique pesticides in total. Figure 1A details the study region and all agricultural pesticide applications reported in 2000, when PEG began.

**Figure 1.**
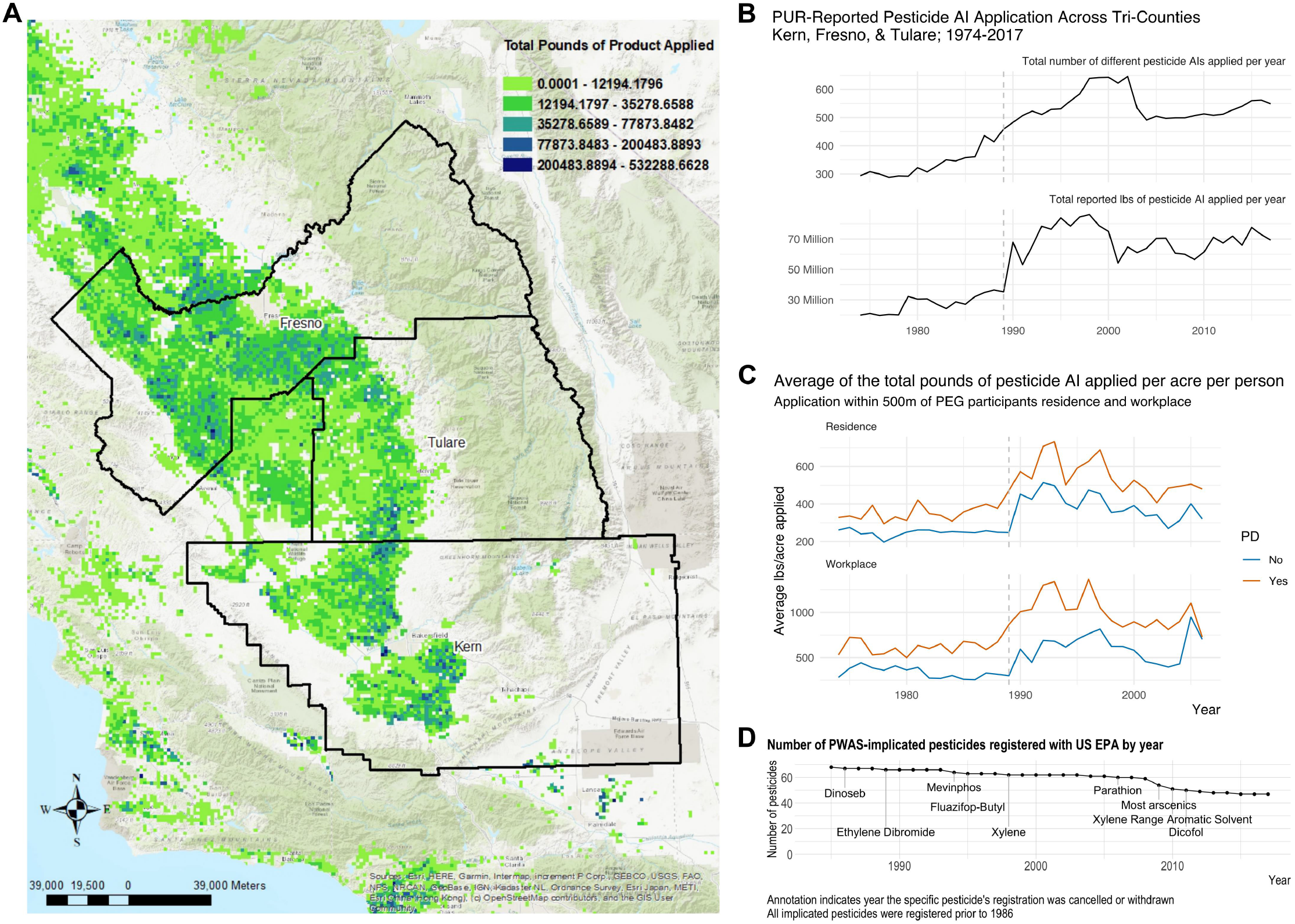
Pesticide Application Descriptives. **(A)** Geography of study region for PEG cohort and total pounds of pesticides applied in the region in 2000. Total pounds of pesticides applied shown by color scale. **(B)** The number of different PUR-reported pesticides applied per year across the three counties and the total reported pounds of pesticide applied per year across the three counties (1974-2017). **(C)** The average total reported pounds of pesticide applied per acre around PEG participants’ residential and workplace addresses per year from 1974-2006 (the mean index year), by PD status. **(D)** Timeline showing the number of PWAS-implicated pesticides that were registered with the US EPA by year. The annotation indicates the year the named pesticide had registration cancelled or withdrawn.

The number of different PUR-reported pesticides applied per year across the three counties since 1974, which ranged from a low of 288 different pesticides applied in 1977 through the high of 646 applied in 2005, is shown in Figure 1B,. Of the 1,355 different pesticides applied from 1974-2017, 722 were applied within the 500m buffer of at least one PEG participant’s residence or workplace. The total reported pounds of pesticide applied per year across the three counties can also be viewed in Figure 1B. The trend in total pounds of pesticide applied since the 1970s increased with time and across the three counties peaking in 1998. There was a notable jump in the total reported pounds applied in 1990, when full reporting to the PUR for all pesticide applications in production agriculture began (see Methods).

On average, the PD patients in the PEG study both lived and worked near commercial agricultural facilities applying more total pounds of pesticide per acre (Figure 1C) than controls (average annual mean difference: 133 more pounds of pesticide applied per acre per year around the patients’ residences versus controls’ and 343 more pounds around workplaces). While summed pounds of applied pesticide across different active ingredients is a crude measure, as the toxicity of each per pound is not necessarily comparable, the trends of total pounds of active ingredient application near addresses closely mimic the trends seen for the entirety of application in the three counties. Of the 722 different active ingredients applied within the study participants’ buffer zone (defined above), PD patients and controls on average lived near the application of 50 (SD=44.4) and 45 (SD=40.9) different pesticides, respectively, during the entire exposure. The mean number near participants’ workplace was 50 (SD=45.4) for patients and 38 (SD=39.2) for controls. Of course, for any given single year, the total number of pesticides was lower (Supplemental Figure 1).

Similar differences were observed in each study wave independently, for men and women separately, and also when limiting to the 288 pesticides with ≥25 exposed participants (Supplemental Table 1). Figures displaying the median values are shown in Supplemental Figure 2.

### Novel pesticides associated with PD in a pesticide-wide association analysis (PWAS)

We assessed each pesticide individually in what we call a PWAS. Of the 722 pesticides with applications near PEG participants’ residence or workplace, we included 288 in our PWAS, according to our criterion of having at least 25 exposed study participants. After limiting to the 288 pesticides, we conducted univariate, unconditional logistic regression to calculate odds ratios (ORs) and 95% confidence intervals (CIs) for PD with each pesticide (n=288). We combined the OR estimates from each study wave and location (residential and occupational addresses) in a fixed effects meta-analysis.

Figure 2A shows a Manhattan plot delineating the statistical significance for each of the 288 tested pesticides, separated by pesticide use type. In total, our PWAS implicated 25 pesticides as associated with PD at a meta-analysis FDR≤0.01 (8.7% of all tested pesticides), another 28 at 0.01<FDR≤0.05 (9.7% of all tested pesticides), and a further 15 at 0.5<FDR<0.10 (5.2% of all tested pesticides) (Figure 2B). Supplemental Table 2 details the OR estimates and 95% CI from the metaanalysis, along with all estimates stratified by study wave and exposure location, for the pesticides with an FDR<0.10. The top five associated pesticides by FDR were sodium chlorate (meta OR=1.24 per SD of log(average pounds/acre), 95% CI=1.14, 1.34, FDR=3.8e-05), dicofol (OR=1.23, 95% CI=1.14, 1.34, FDR=4.2e-05), prometryn (OR=1.24, 95% CI=1.14, 1.34, FDR=6.0e-05), methomyl (OR=1.20, 95% CI=1.11, 1.30, FDR=3.8e-04), and xylene range aromatic solvent (OR=1.20, 95% CI=1.10, 1.30, FDR=8.8e-04). These pesticides showed consistent risk profiles for both residence and workplace address locations and were found to be associated with PD in both PEG1 and PEG2 for at least one of the two locations (Supplemental Figure 3). Exposure descriptive statistics, including exposure prevalence, and the risk estimates for pesticides associated with PD at p<0.10 can be found in Supplemental Tables 3 and 4.

**Figure 2.**
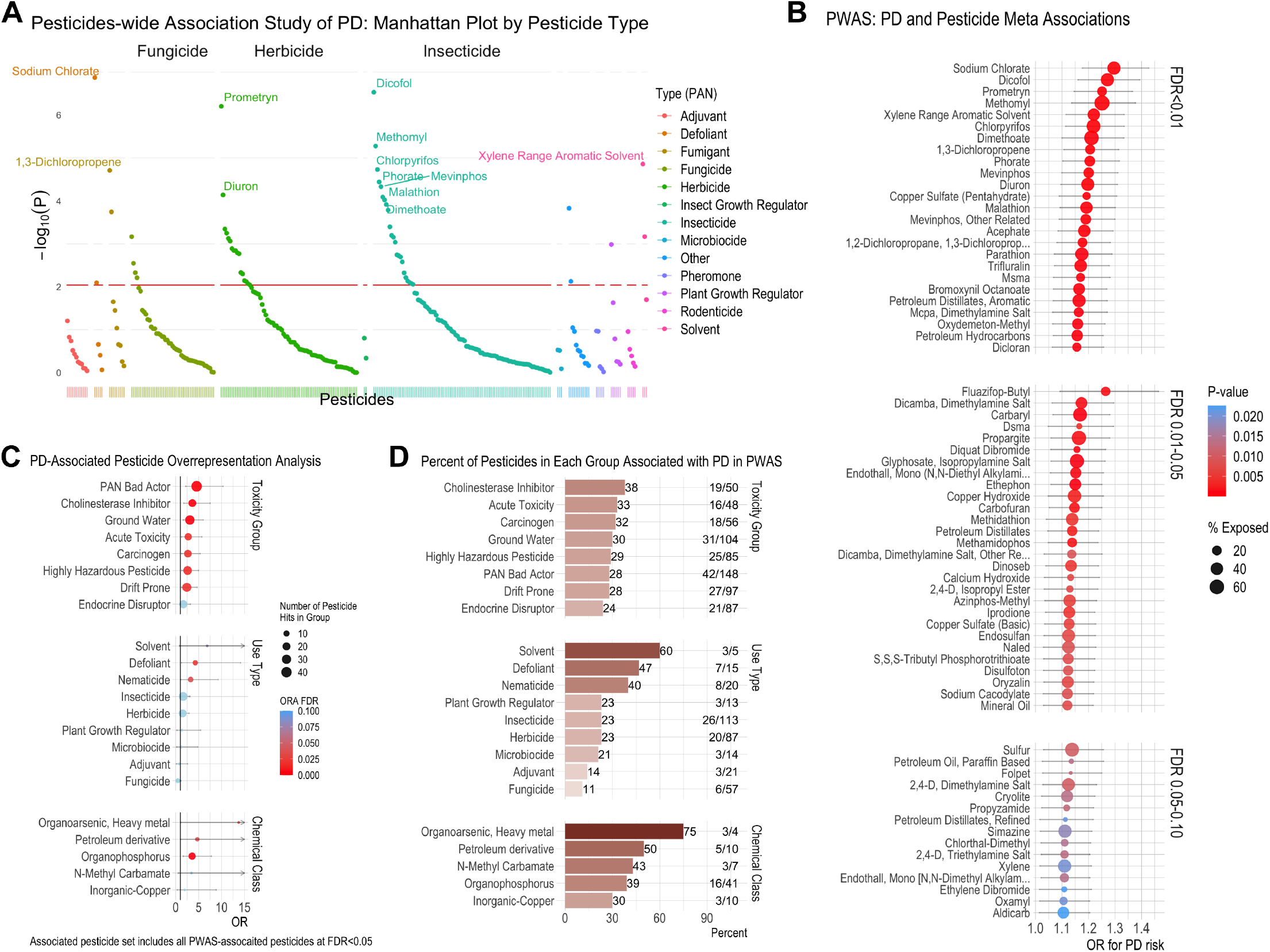
PWAS analysis. **(A)** Manhattan plot detailing the negative log p-value from the meta-analysis for all n=288 pesticides tested for association with PD. **(B)** Dot plot displaying the odds ratio (OR) and 95% CI from the metaanalysis for all pesticides with an FDR<0.10. **(C)** Results of overrepresentation analysis to test for overrepresentation of pesticide groups (toxicity groups, chemical classes, and use types) in the set of PWAS PD-associated pesticides relative to all pesticides we assessed. **(D)** Bar graph indicating the total number of pesticides tested in the PWAS in each group, the number of pesticides in each group PWAS-associated with PD, and the percent. For example, 50 cholinesterase inhibitor pesticides were assessed for association with PD, 19 (38%) were associated with PD in the PWAS.

Regulatory and toxicity information for the PWAS-implicated pesticides is shown in Supplemental Table 5. This provides information on current registration and usage in California, the US, and the European Union (EU) as well as toxicity designations. Eighteen of the 25 most strongly PD-associated pesticides (FDR<0.01) with use during our exposure window are actively registered with the EPA allowing for current use in the US (43 of 68 pesticides at FDR<0.10 are currently registered, Figure 1D), while only 6 are allowed for use in the EU. Of these 25 pesticides, 21 are considered ‘bad actors’ by the Pesticide Action Network (PAN) as 9 have been deemed carcinogens (7 more are possible carcinogens), 6 developmental or reproductive toxins, 10 cholinesterase inhibitors, 3 known groundwater contaminants (13 more are possible groundwater contaminants), and 8 have high acute toxicity. In fact, of the 53 pesticides with an FDR<0.05, 43 have been designated ‘bad actors’ by PAN.

We used overrepresentation analysis (ORA), which is commonly applied to evaluate gene set overrepresentation, to test for enrichment of pesticide groups (toxicity groups, chemical classes, and use types) in the set of PD-associated pesticides relative to all pesticides we assessed. We linked each of the 288 pesticides included in the PWAS to chemical, regulatory, and toxicity information using publicly available databases. We assessed overrepresentation of the PD-associated pesticide set (PWAS FDR<0.05, n=53 pesticides) relative to all pesticides assessed (n=286 pesticides with group classifications) for all groupings (toxicity classifications, use types, and chemical classes) with at least 3 associated pesticides in the group (n=22 groups). The results of the ORA are shown in Figure 2C and Supplemental Table 6, while Figure 2D shows the percent of pesticides associated with PD in each group (i.e. 48 of all 286 tested pesticides are considered acutely toxic, 16 of these 48 were associated with PD at FDR<0.05 (33%)).

Based on the ORA, nearly every toxicity classing was overrepresented in the set of PD-associated pesticides relative to all pesticides assessed. This indicates there were more PD-associated pesticides with these toxicity classifications than expected based on the group’s distributions among all assessed pesticides. For example, 16 of the 53 pesticides (30%) associated with PD are considered acutely toxic, while only 17% of all tested pesticides are considered acutely toxic. Using a Fisher’s exact test and odds ratio (OR=2.70, 95% CI=1.25, 5.69, FDR=0.02), we found that the odds of being among the PD-associated pesticides is 2.7-fold higher for the acutely toxic pesticides versus the non-acutely toxic pesticides. The following toxicity groupings were also significantly overrepresented in the PD-associated pesticide set, Highly Hazardous Pesticides: OR=2.56 (95% CI=1.32, 4.97); Cholinesterase Inhibitors: OR=3.62 (95% CI=1.73, 7.50); and Carcinogens: OR=2.63 (95% CI=1.26, 5.36). Furthermore, the odds of being among the PD-associated pesticides was also 2 to 3-fold higher for pesticides that contaminate groundwater (OR=3.08 (95% CI=1.60, 5.99) and for those that are highly prone to drift (OR=2.41 (95% CI=1.26, 4.64) (Supplemental Table 6).

Several pesticide chemical classes and use types were also overrepresented, though small numbers limited statistical power for most groups. These included the defoliant (OR=4.25, 95% CI=1.25, 14.17), nematicide, and solvent use types. They also included organophosphorus, heavy metal organoarsenic, and petroleum derivative chemical classes. For example, while only four heavy metal, organoarsenic pesticides were tested for association with PD in our PWAS, three (or 75%) were associated with PD (see Figure 2D).

### Multiple PWAS-identified pesticides are toxic to iPSC-derived dopaminergic neurons

PD is increasingly thought of as a multi-system disease involving both neuronal and extraneuronal tissues [19, 20]. Pesticides could thus confer increased PD risk in many different ways. Some, however, are presumably directly toxic to mDA neurons. We tested this directly in iPSC-derived mDA neurons derived from a patient with PD who harbored a pathologic triplication at the α-syn-encoding *SNCA* locus[13]. This line, derived from a member of the so-called Iowa kindred, was selected because it over-expresses wild-type α-syn and reflects an extreme form of the pathology associated with idiopathic PD, namely the accumulation in neurons of wild-type α-syn. Neurons derived from this iPSC line exhibit PD-relevant phenotypes [21–25]. We engineered these iPSCs to express a fluorescent reporter at the tyrosine hydroxylase (TH) locus to overcome the heterogeneity of mDA differentiation. This TH tdtomato reporter was targeted via CRISPR-Cas9 and confirmed via PCR and Sanger sequencing (Figure 3A, B, C, [15]). Colocalization of endogenous THtdtomato signal with anti-TH-labeled cells was confirmed. Localization was consistent with the expression pattern observed in mDA neurons derived from embryonic stem (ES) cell reporter lines (Figure 3F, [15]. This reagent enabled us to selectively test the effects of pesticides on mDA neurons rather than other contaminating cell types that variably co-exist in mDA cultures differentiated from iPSC.

**Figure 3.**
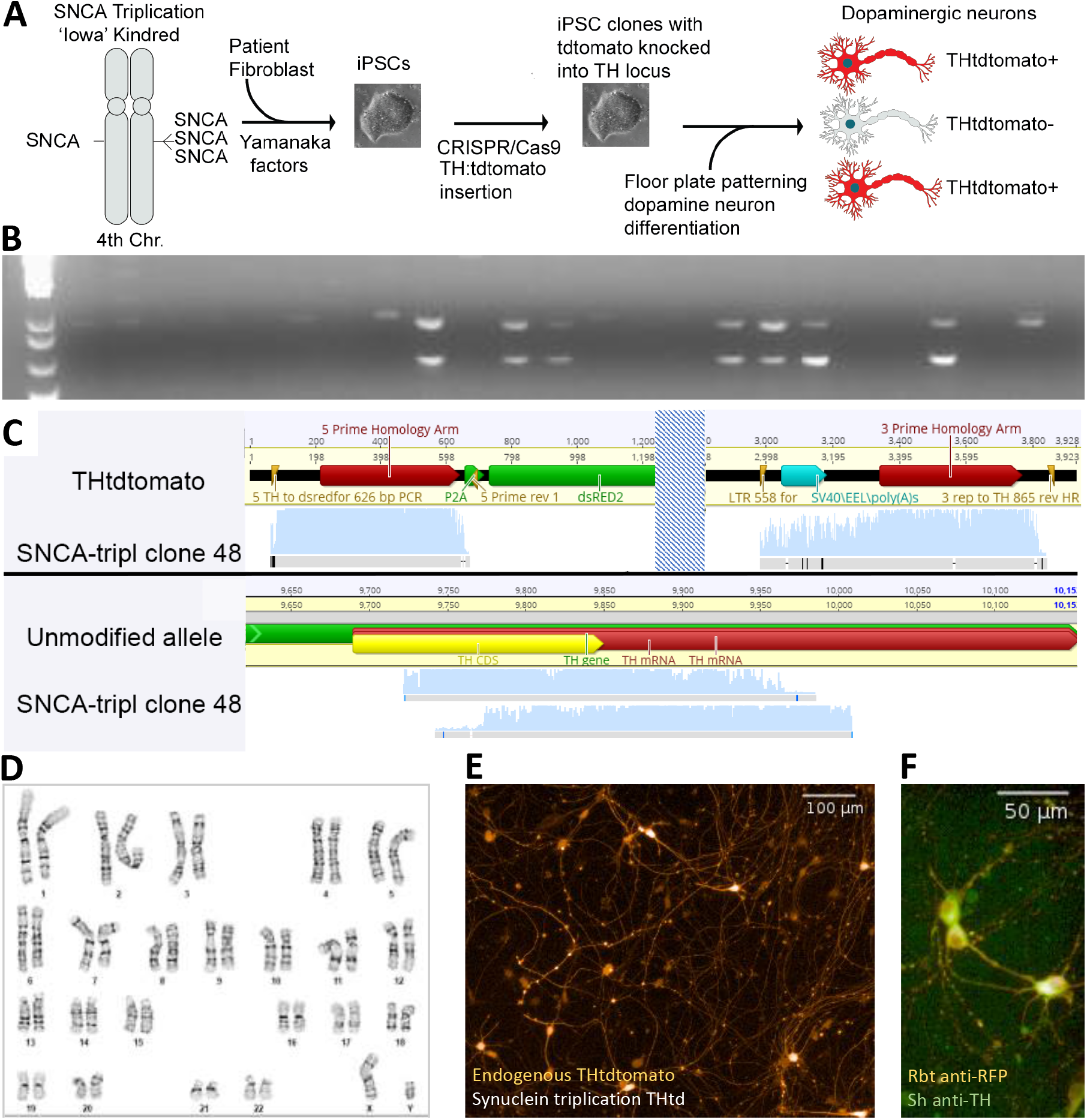
Synuclein triplication THtdtomato reporter generation. **(A)** Schematic demonstrating iPSC source, generation, modification, and differentiation with tdtomato reporter permitting identification and isolation of dopaminergic neurons. Previously described iPSC line derived from a patient with Parkinson’s Disease caused by triplication of the alpha synuclein locus resulting in four copies of the gene. iPSC line was then modified with tyrosine hydroxylase:tdtomato reporter [15]. Adapted from Hallaci et al *in revision*. **(B-F).** Quality control and validation of synuclein triplication THtdtomato reporter line including 5’ and 3’ PCR products to confirm proper insertion. (**B**) Agarose gel stained with ethidium bromide to demonstrate examples of seven clones that contain the expected PCR products (626 bp product confirmed proper insertion of the 5’ end of the reporter construct and an 878bp product confirming proper insertion at the 3’ end). PCR reactions run separately but combined into the same wells for each clone to visualize clones passing and failing PCR quality control. A subset of clones have a single larger band and these are excluded from further testing. (**C**) Sanger sequencing of PCR products in (B) confirming correct insertion of tdtomato cassette. (**D**) G-banded karyotype (performed by WiCell) confirms normal karyotype in modified clone. (**E**) Example of live imaging of endogenous THtdtomato fluorescence at 10x. (**F**) Immunofluorescence co-localization of Rabbit anti-RFP and Sheep anti-tyrosine hydroxylase visualized with Alexa Fluor 546 donkey antirabbit and Alexa Fluor 488 donkey anti-sheep, respectively.

Forty-three of the pesticides identified in the above PWAS analysis were obtained and resuspended in DMSO, water, or ethanol based on available solubility data. A four-point dose curve protocol was used spanning a range of pesticide concentrations comparable to what has been used in previous toxicity assays utilizing the Tox21 compounds on multiple human cell lines including neural lineage cells [26, 27]. THtdtomato-positive neurons were fluorescence activated cell sorted (FACS) (Supplemental Figure 3) and used to test for sensitivity to the pesticides in a live imaging survival assay. Quantitation of THtdtomato-positive mDA neurons was performed eleven days after initial treatment (Supplemental Figure 4). A dynamic range was established between negative controls (DMSO, water, DMSO+Ethanol) and positive pesticide controls. The Z-prime score for the DMSO negative control and rotenone pesticide positive control was 0.549. Ziram, described previously as toxic to mDA neurons [28], served as an additional positive control. Exposure to 10 pesticides led to cell death >3 standard deviations greater than the DMSO control mean at a concentration of 30μM: propargite, copper sulfate (basic and pentahydrate), dicofol, folpet, naled, endothall, trifluralin, endosulfan, and diquat dibromide (Figure 4). For example, propargite treatment produced extensive DA neuron death and degeneration of neurites (Figure 4 *right upper* and *right lower*). Dose-response curves extending below the screening concentration indicated that propargite was the most toxic, followed by diquat dibromide, folpet, and naled. All were toxic at 6μM as well (Figure 5). The large standard deviation seen for dicofol was due to an outlier. Subsequent repeat dose-response curve confirmed it is toxic at concentrations of 9.5μM and greater (Supplemental Figure 5). Information about pesticide use type, chemical class, regulation status, and prior toxicity classifications for the 10 pesticides which produced substantial cell death is included in Supplemental Table 7. In summary, our field-to-bench platform identified compounds that were associated with an increased risk of PD and were directly toxic to mDA neurons. These compounds encompassed a range of use types (four insecticides, three herbicides, and two fungicides); were structurally distinct; and do not have an overlapping prior toxicity classification, such as acute toxicity.

**Figure 4.**
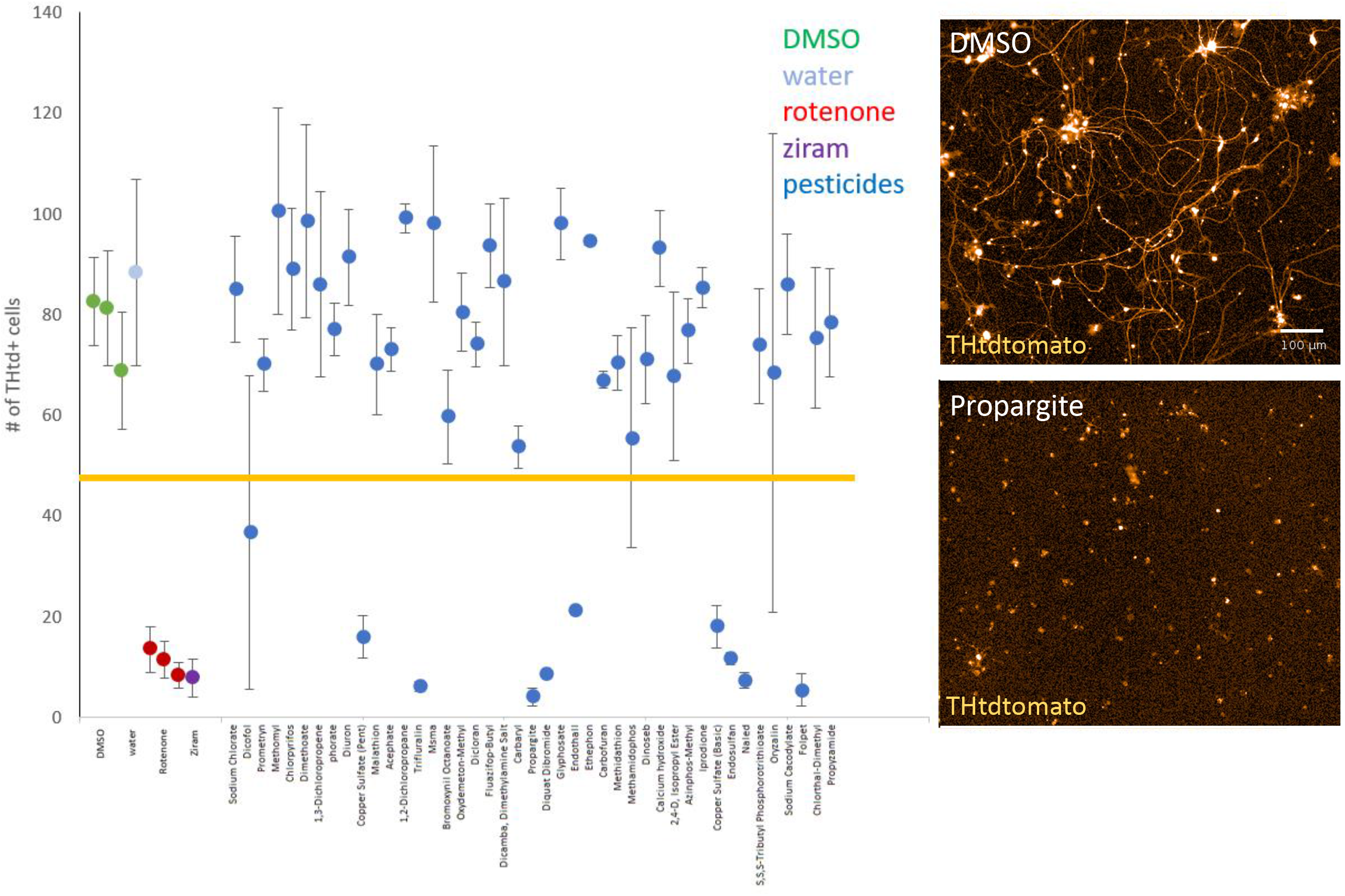
PWAS pesticides are directly toxic to mDA neurons. Scatter plot with the number of THtdtomato+ cells measured by live imaging analysis 11 days after the first treatment. DMSO controls (green data points) were present on each assay plate. Water control (light blue) was present on the assay plate containing primarily water-soluble pesticides. Rotenone (red data points) and ziram (purple data point) were used as positive controls. Blue data points represent the different pesticides from the PWAS study. Error bars are two standard deviations (one above, one below). Large standard deviations were observed for pesticides which had a single outlier out of three technical replicates (dicofol, methamidophos, oryzalin). Horizontal line denotes three standard deviations below DMSO mean. Right upper image is a 10x magnification live image of a DMSO control well. Right lower image is from a propargite well. Scale bar = 100μM

**Figure 5.**
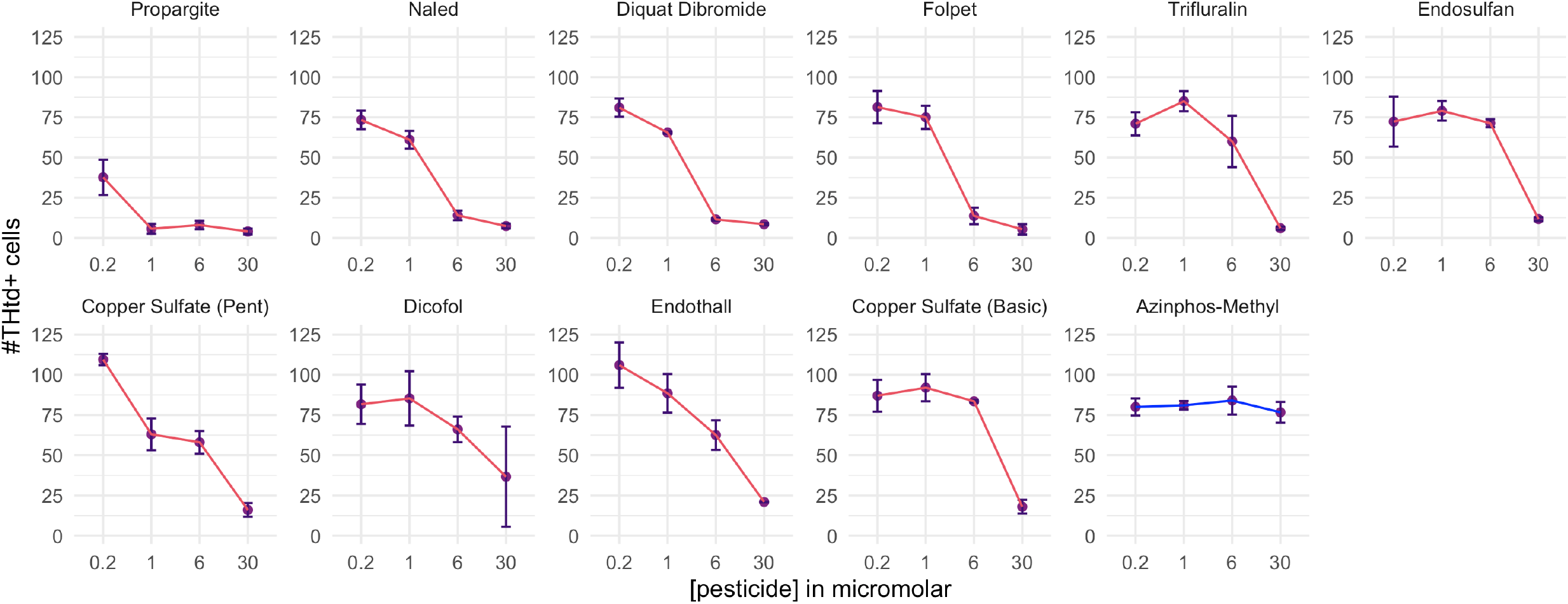
Dose response curves of most toxic pesticides. Four concentration dose curves of PWAS toxicants producing death in synuclein triplication THtdtomato sorted neurons. Cell numbers measured by high content imaging of live cultures 11 days after first treatment. Error bars are two standard deviations (one above, one below).

### PWAS-identified pesticides cluster based on correlation of exposures

Pesticides are not applied in isolation. Combinations of different pesticides are regularly applied to the same fields within the same season, year after year. Some are applied as part of the same product, while others as combinations of different products. We thus investigated how exposure among the PWAS-implicated pesticides was correlated. Further, we asked how the pesticides directly toxic to mDA neurons related to other PWAS-implicated pesticides, either those that did not produce significant mDA death or those that could not be tested. Finally, we used real-world exposure clusters to motivate combinatorial assessment of pesticides, and potential synergistic toxicity, in our mDA neuron assay.

Figure 6A shows a heatmap detailing the correlation between all PWAS-implicated pesticides (FDR<0.10, n=68 pesticides). The heatmap, which shows broad patterns of correlation, indicates multiple groups of highly correlated pesticides. The complete pairwise correlation tables are detailed in Supplemental Tables 8 and 9. In order to specifically assess how the mDA toxic pesticides correlated with the other PWAS-implicated pesticides, we used a data-driven integration and network analysis approach to assess correlations across two layers (Figure 6B). The two layers were, first, the set of mDA toxic pesticides, designated in the correlation circle as teal highlighted diamonds, and, second, the set of all other PWAS-implicated pesticides, shown as circles. All correlations between layers at R>0.45 are shown in the circle, while correlations within layers are not shown. The size of the shapes in the correlation circle (diamonds and circles) were determined by the PWAS FDR, so pesticides that were more strongly associated with PD in the PWAS are represented by larger sized shapes. In contrast, the color of the shapes reflects the density of the connections (i.e. correlations at R>0.45) made by that specific pesticide with others. Pesticides with a darker color are correlated with more pesticides, and arrangement around the circle is ordered from those with the most correlations (dicofol, darkest color) to the least (petroleum hydrocarbons, lightest color). Dicofol, for example, resulted in significant mDA cell death in the iPSC-model and is therefore shown as a teal highlighted diamond. It was also both (1) the most statistically significant mDA toxic pesticide in the PD PWAS (FDR=4.2e-05) and therefore shown as the largest diamond, and (2) correlated above R>0.45 with the most other PWAS-implicated pesticides (n=24 pesticides), and therefore shown as the darkest color. In the correlation wheel, each mDA toxic pesticide can be assessed in this manner.

**Figure 6.**
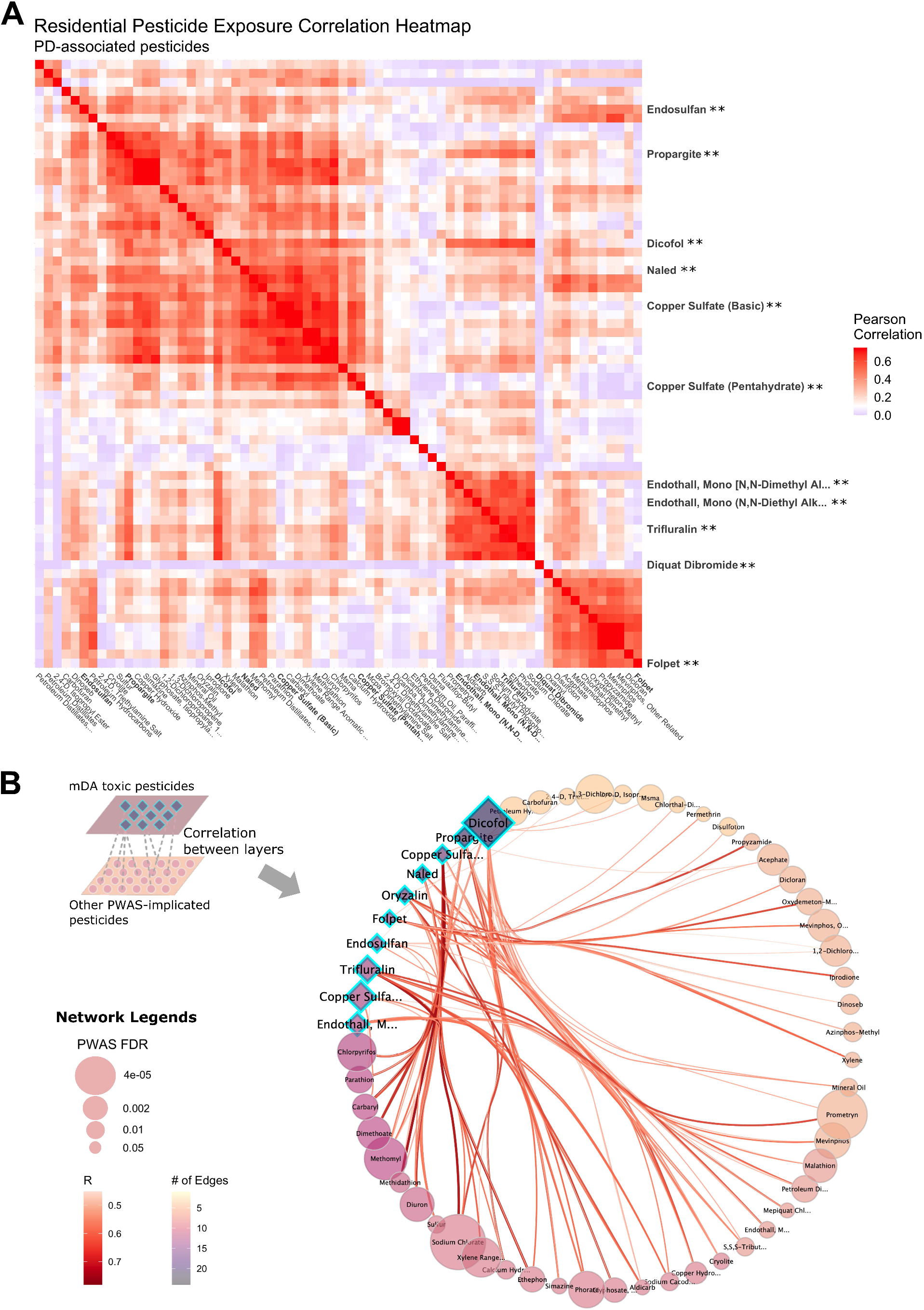
Pesticide Exposure Correlations. **(A)** Correlation heatmap indicating the pairwise Pearson correlation coefficient for 68 PWAS-implicated pesticides (FDR<0.10), using residential address-based exposures. The pesticides which produced significant mDA neuron death in the iPSC-model are highlighted on the y-axis. No pesticides were significantly negatively correlated, thus the blue color represents null (R=0) correlation to the dark red (R=1.0) **(B)** Correlation wheel showing the pesticide exposure correlations across two layers: the set of mDA toxic pesticides, which are designated as teal highlighted diamonds, and the set of all other PWAS-implicated pesticides, shown as circles. Correlations between layers at R>0.45 are shown in the circle, correlations within layers are not shown. The shape (diamonds and circles) size was determined by the PWAS FDR, indicating pesticides that were more strongly associated with PD in the PWAS are bigger. The color of the shape was determined by the number of pesticides in the opposite layer which the pesticide correlates with at R>0.45. Pesticides with a darker color are correlated with more pesticides, and the circle is ordered from those with the most correlations (dicofol, darkest color) to least (petroleum hydrocarbons, lightest color).

Overall, this analysis showed that exposure to the mDA neurotoxic pesticides associated with PD was highly correlated with exposure to other PD PWAS-associated pesticides. Specifically, exposure to 75% of the pesticides associated with PD in the PWAS that either did not result in significant mDA cell death or could not be tested was correlated with exposure to at least one pesticide that did result in significant mDA cell death (R>0.43). Put another way, exposure to the majority of the pesticides tied to PD in our PWAS, including those that did not directly lead to mDA neuron toxicity in our assay, was highly correlated with exposure to a pesticide that did exhibit such toxicity. Some compounds form correlation network hubs, consisting of the most interconnected to other pesticides. Dicofol emerged as the strongest hub (highest closeness centrality and highest number of edges), followed by propargite and methomyl.

Next, we defined correlation clusters representing real-world co-exposure relying on an unsupervised, hierarchical clustering approach (Figure 7A). We separated all PWAS-implicated pesticides into groups based on a correlation coefficient R>0.45. In total, we detected 25 different pesticide clusters using residential-based exposure. These are shown as different colors in the dendrogram (Figure 7A) and detailed in Supplemental Tables 10 and 11 (residential and workplace exposure clustering). Some of the clusters consisted of a single pesticide, meaning these pesticides did not correlate above 0.45 with any other pesticide. The other clusters represent co-exposure profiles of interest for further *in vitro* testing.

**Figure 7.**
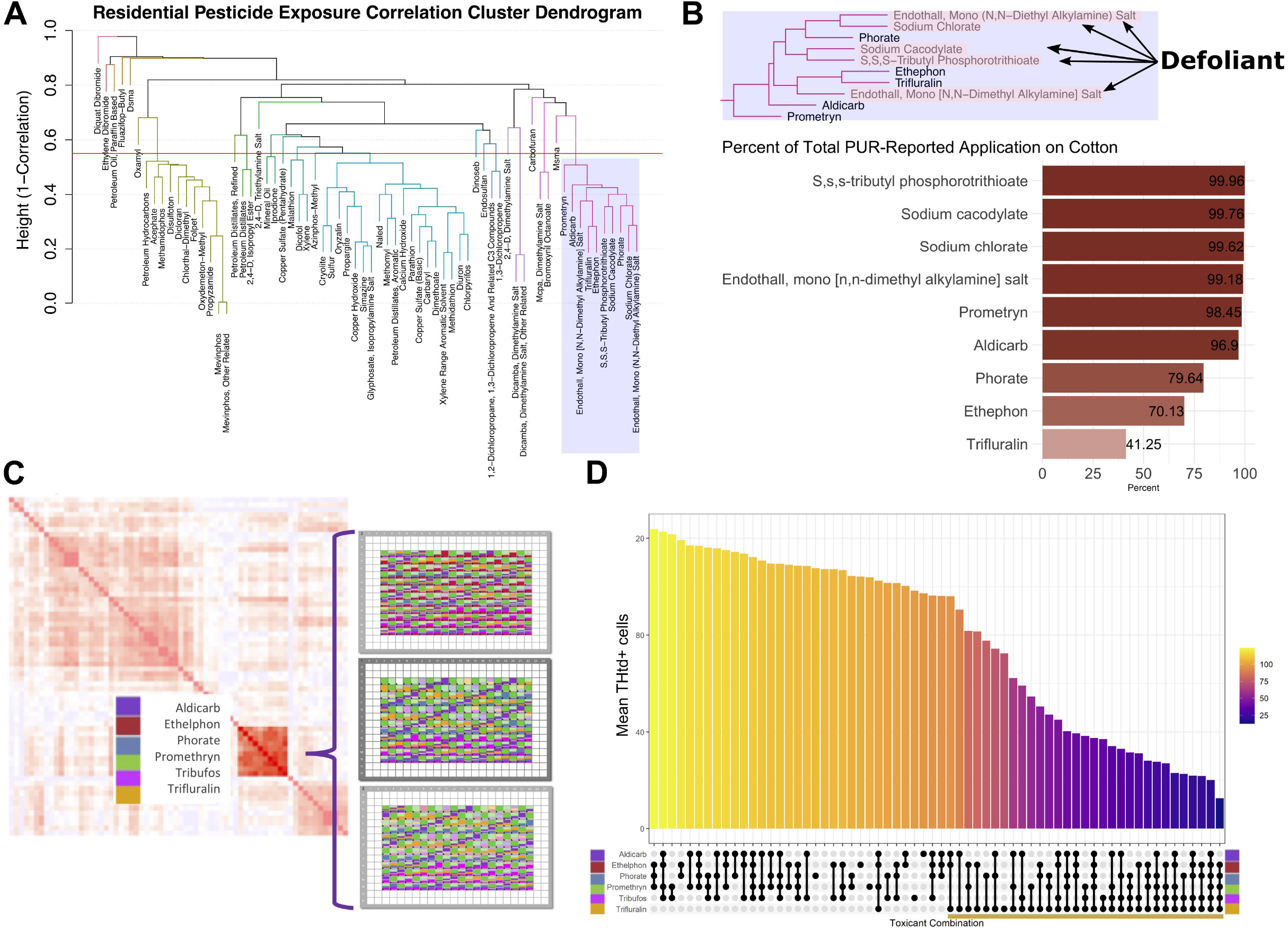
Synergy of pesticides in the Cotton Cluster. **(A)** Cluster dendrogram from hierarchical clustering of the PWAS-pesticides using residential address-based exposures (clusters cut at height=0.55), to identify groups of highly correlated pesticides for coexposure analysis. **(B)** Cotton cluster: particular cluster of interest identified, as it includes two of the three most significantly associated pesticides in the PWAS, based on FDR (sodium chlorate and prometryn) and half of the cluster have a use type of defoliant. The bar graph shows the proportion of all agricultural application records in the tri-county study area used on cotton, for each pesticide in the cluster. For example, 99.96% of the reported S,S,S-tributyl phosphorotrithioate (aka tribufos) applications were on cotton. **(C)** Schematic outlining how pesticides from a single co-exposure cluster (cotton cluster) were recombined in all possible combinations of six pesticides using an HP Digital dispenser on sorted dopaminergic neurons plated into a 384 well format, similar to the survival assay described in Figure 4. **(D)** An upset plot was used to sort and represent the most toxic and least toxic combinations. Y-axis shows number of THtdtomato+ neurons at day 11 following treatment. Ball and stick connections along the X-axis indicate co-treatments with a ball indicating treatment with a given pesticide. DMSO control condition is depicted by the x-value lacking any ball and stick marker.

We isolated one cluster of particular interest with clear co-application patterns that also contained several of the most statistically significant PD PWAS-pesticides, including the two of the top three hits, sodium chlorate and prometryn (Figure 7B, Cluster 7 in the supplemental tables). Half of this particular cluster was defoliants, a unique type of pesticide which causes leaves to fall off plants. Defoliants are applied in agriculture almost exclusively on cotton. We confirmed this by aggregating all PUR records in the tri-county area over the study period (~5.9 million records) and assessing the proportion of the applications for each pesticide on cotton (Figure 7B). For example, we found 99.96% of the reported S,S,S-tributyl phosphorotrithioate (tribufos) applications were on cotton, 99.76% of sodium cacodylate, and 99.6% of sodium chlorate. Given the clear patterns of real-world co-exposure due to proximity to cotton agriculture, strength of the individual PWAS associations, and manageable cluster size for *in vitro* combinatorial testing, we selected members of this “cotton cluster” for further coexposure experiments *in vitro* on mDA neurons.

### *In vitro* synergy testing identifies pesticides with most potent direct effects on dopaminergic neurons from the cotton cluster

Co-exposures can be modeled *in vitro* via concurrent treatment to directly compare the consequences of multiple direct co-exposures or via iterative combinations of the co-exposure cluster. We hypothesized that pesticides with the most relevance to PD risk would be likely to cause mDA neuron cell death on their own, but that toxicity could be enhanced by co-exposure to other pesticides commonly applied to the same fields (i.e. from the same exposure cluster). To test effects of coexposure to the cotton cluster, we performed a survival assay on sorted mDA neurons using combinations of pesticides in this cluster (Figure 7C). The schematic in Figure 7C illustrates how pesticides from a single co-exposure cluster (shown in more detail in Figure 6A) were combined in an exposure matrix, such that each pesticide was added to mDA neurons in every possible combination with the other pesticides chosen for the experiment. We used a live imaging survival assay and treatment paradigm similar to that used for data presented in Figure 4. A digital dispenser (Hewlett Packard) was used to design and dispense pesticides for the full set of combinations.

A 10μM dose was chosen for this assay given that this represents an intermediate dose for one pesticide in the group that was toxic at 30 μM and could permit detection of additive or synergistic effects. Number of surviving neurons after treatment were then arranged in descending order to determine the most toxic combinations. (Figure 7D). We performed four independent biological replicates and the average data are presented. P-values for individual pairwise comparison with adjustment for multiple testing provided in Supplemental Table 13. These data indicate that combinations involving trifluralin cause more dopaminergic neuron cell death (gold bar) and potential synergy with tribufos (S,S,S-tributyl phosphorotrithioate) as this combination results in the most mDA neuron cell death of all the pairwise combinations. Trifluralin alone at 10μM produced a 32% decrease in mDA neurons compared to DMSO, while tribufos produced an 8% decrease (neither statistically significant in this assay), but in combination they produced a 65% decrease compared to DMSO control and were significantly different from the individual treatments at p= 0.003 for the comparison to tribufos alone and p = 0.048 for the comparison to trifluralin alone. Performance in this assay showed no clear correlation to the odds ratio for PD risk or FDR cutoff level described in Figure 2B.

### Trifluralin reduces spare capacity of mitochondria in mixed dopaminergic neuron cultures

Our field-to-bench paradigm pinpointed trifluralin as a pesticide associated with PD in the PWAS screen that was both toxic to mDA neurons *in vitro* (Figure 4) and interacted synergistically with other co-applied pesticides in the cotton cluster. To more deeply investigate potential mechanisms by which trifluralin causes mDA neuron cell death, we plated cultures enriched for mDA neurons at day 35 of differentiation and assessed effects on mitochondrial function. We chose mitochondrial function because extensive literature has documented the selective vulnerability of these neurons to mitochondrial dysfunction. Specifically, we quantified relative mitochondrial subunit abundance, mitochondrial respiration, and function of the oxidative phosphorylation pathway in mDA neuron cultures. Cultures were maintained for at least 3 weeks after plating prior to analysis. This prolonged culture period promoted complex neurite arborization and additional neuronal maturation. We measured mitochondrial subunit abundance after 24-hour exposure to 30μM trifluralin in 3 biological replicates and demonstrated a 50% reduction of Complex I (NDUFB8) and Complex IV (COX II) when compared to DMSO treated controls (Figure 8A). There was no significant reduction or increase of the expression of other complexes in the mitochondrial respiratory chain.

**Figure 8.**
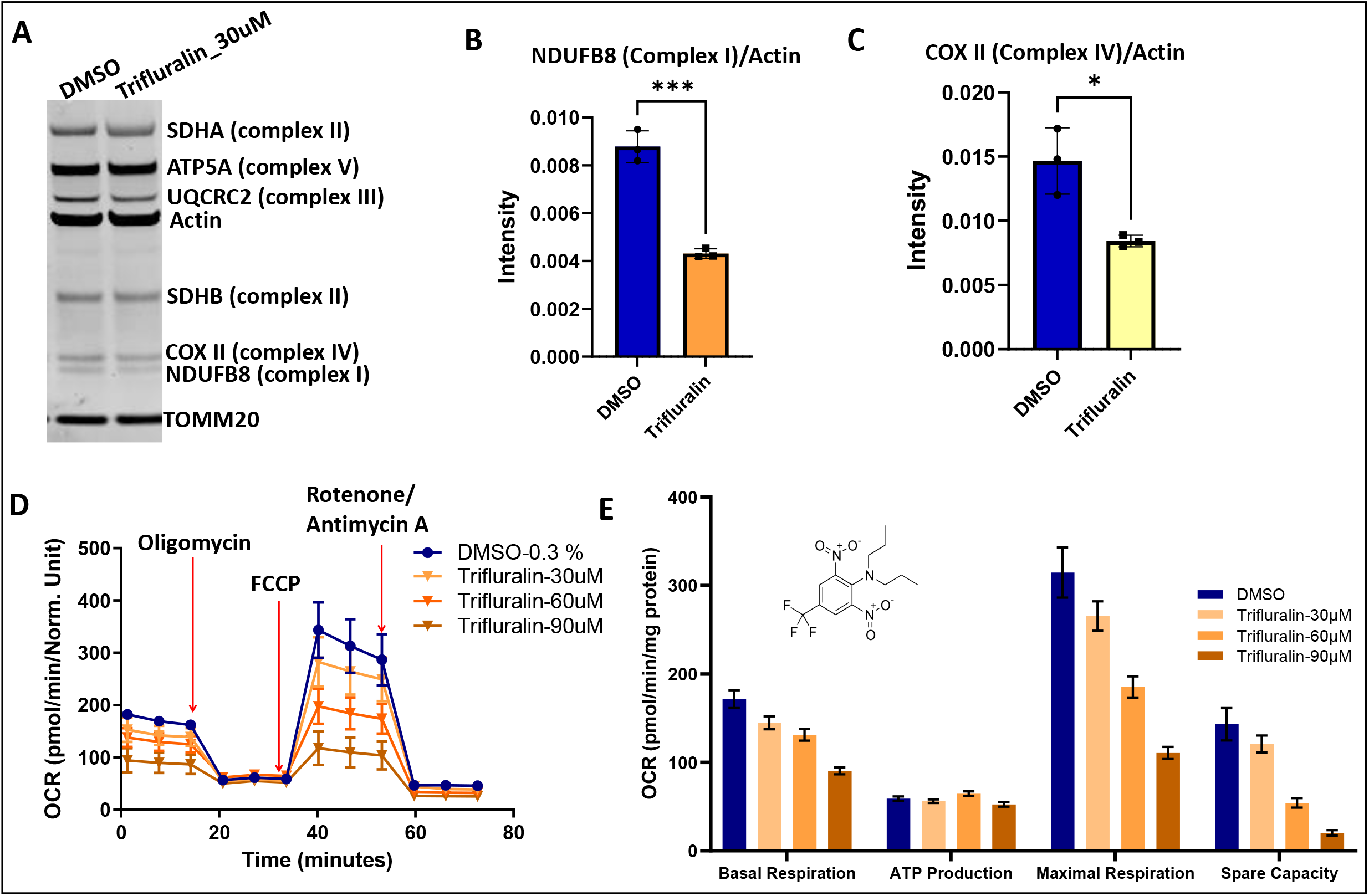
Trifluralin alters mitochondrial subunit abundance and oxygen consumption rate. Effect of trifluralin in mitochondrial subunit and MitoStress assays. (A)Western blot for respiratory chain complexes for differentiated SNCA-triplication neurons at DIV 65, exposed to 0.3 % DMSO and 30 μM trifluralin for 24hr. (**B, C**) Quantification of Complex I (**B**), p=0.0004 and Complex IV (**C**), p=0.0145 from blot in (**A**). (**D**) Measurement of Oxygen Consumption Rate curves on Agilent Seahorse XFCell Mito stress assay for the dose response effect of DMSO (0.3%) and trifluralin (30 μM, 60 μM and 90 μM) on SNCA-triplication dipg18fferentiated neurons at DIV 65 and after 6hr exposure. (**E**) Metabolic parameters calculated from D.

For assessment of mitochondrial function, we used the well-established Seahorse assay. Briefly, this assay involves sequential addition of mitochondrial complex inhibitors (Oligomycin, FCCP and Rotenone/Antimycin-A) and measurement of media acidification and oxygen consumption rate. The effects of trifluralin were assessed in a dose response manner, using 0.3% DMSO and trifluralin at 30 μM, 60 μM and 90 μM (Figure 8D). We then calculated mitochondrial respiration parameters (basal respiration, ATP production, maximal respiration and spare capacity) and found, with the exception of ATP production, all were decreased with exposure to trifluralin at the concentrations used in the assay. The respiratory capacity of differentiated neurons was reduced to 14% when cells were treated with trifluralin at 90 μM, 38% at 60 μM, and 84% at 30 μM trifluralin compared to the treatment with DMSO (Figure 8E). The mitochondrial subunit assay highlights an effect on Complex I and Complex IV when differentiated neurons are treated with trifluralin (Figure 8). The effect of trifluralin on cellular respiration is dose and time-dependent (Figure 8D-F)

We also exposed mDA neurons to ziram. Ziram has been implicated in PD previously and its toxicity has been tied to an inhibitory effect on the E1 ligase in mDA cultures without any described mitochondrial phenotypes [28] (Supplementary Figure 6A). Indeed, we found that ziram exposure for 2 or 6 hours at a dose greater than the LD50 from survival assays (300nM) did not significantly alter the oxygen consumption rate (Supplementary Figure 6C) or the mitochondrial reserve capacity (14% increase in 6 hr exposure compared to DMSO) (Supplementary Figure 6B). Treatment with trifluralin for the same amount of time at a roughly equivalently toxic dose of 30μM (Supplementary Figure 6F) led to a decrease in oxygen consumption rate consistent with a reduction in mitochondrial reserve capacity to 70% compared to DMSO (Supplementary Figure 6D-E). These data implicate mitochondrial dysfunction specifically in trifluralin-mediated mDA neuron cell death. More generally, our data highlight that pesticides toxic to mDA neurons exhibit distinct mechanisms of toxicity.

## DISCUSSION

Pesticides have long been linked to Parkinson’s disease etiology. However, most pesticides used in agriculture have not been assessed for potential influence on PD. Therefore, we established a field-to-bench paradigm which combined two distinct approaches: 1) broad epidemiologic screening of hundreds of pesticides for association with PD; and 2) *in vitro* evaluation of PD-associated pesticides in mDA neurons derived from patient iPSCs. This approach permitted testing of epidemiologic hits for direct effects on dopaminergic neurons to better identify and classify PD-relevant pesticides.

Our record-based exposure assessment, which takes advantage of the California Pesticide Use Report system detailing 50 years of agricultural pesticide applications, allowed us to screen nearly 300 specific pesticides for association with PD in an untargeted manner. We implicated 68 pesticides with PD using this approach. Many of these present novel targets for investigation, as they have not been implicated as relevant to PD etiology previously. However, this screening in population-based data does not suggest that every pesticide that screened positive causes PD. Equally, we would not conclude that those which failed to screen positive using our PWAS approach, based on poundage of toxicant applied near residences or workplaces, are not relevant to PD etiology. Instead, this approach was a first step to prioritize agents and mixtures of agents for more in-depth research.

To identify whether these pesticides might contribute to PD through direct effects on mDA neurons, we coupled our PWAS screen to systematic analysis of the hits in human mDA neurons derived from iPSCs of a patient with a sensitizing PD mutation (a triplication of the wild-type *SNCA* gene). Of the 43 pesticides we tested *in vitro*, 10 resulted in substantial mDA neuron cell death at 30uM. The identified toxicity in these neurons represents a unique and novel common characteristic of these pesticides--they do not have another previously described shared characteristic or property. We would posit that patient-derived stem cell models may help direct future mechanistic study and provide novel disease-specific pesticide classifications. This represents an extension of an approach that others have taken with individual tool-compound pesticides[4, 29]. Interestingly, our experiments identified propargite and confirmed previous work with this pesticide in mDA neurons derived from a different iPSC line and utilizing a different assay [29]. This orthogonal confirmation indicates the platform may be robust. Personalized *in vitro* disease modeling similar to what we have presented here could provide substantial insight into gene-environment interactions in PD. While considerable focus in the last two decades in PD research has been on genetics, the majority of the risk for PD is likely environmental or tied to complex gene-environment interactions. Our models will now enable detailed investigation of these interactions in distinct genetic backgrounds and also in populations of iPSC-derived neurons grown in a pooled “village” format [30].

The iPSC line used here was derived from a patient with triplication of the wild type α-syn locus. This genetic background is pertinent as it mimics the accumulation of wild-type α-syn that accompanies all cases of so-called sporadic late-onset PD. As described above, neurons made from this iPSC line exhibit PD-relevant phenotypes. These documented phenotypes increased confidence that more subtle sensitivity to pesticides could be amplified by the use of these neurons rather than neurons from a healthier, wild type line. Perhaps most importantly, these cells capture human biology that may differ in critical ways from rodent or other human cell lines: a recent study showed that human mDA neurons from both familial and sporadic PD exhibit unique oxidative biology, including dopamine oxidation, that is not recapitulated faithfully in mouse models [31]. Our current work has detailed specific vulnerability of these mDA neurons to perturbations of oxidative metabolism by trifluralin. Future work will clarify the extent to which this mDA toxicity represents a unique susceptibility of the triplication cell line or is generalizable to other PD cell lines. Follow up investigation will benefit from the use of isogenic lines to specifically assess gene-environment interactions, as has been done for paraquat in a previous study of the A53T α-synuclein mutation[4].

Most people living in agricultural communities are co-exposed to multiple pesticides, including as a result of co-application of different pesticides with the same seasonal application patterns year after year. An important use of our platform is to screen both epidemiologically and experimentally for pesticide co-exposures, and to understand mechanisms of synergistic interactions. For the majority of the pesticides we associated with PD in our PWAS, even if the pesticide itself did not result in significant mDA cell death in our iPSC-model, its exposure levels correlated relatively highly with a pesticide that did. This finding presents some interesting interpretations for the lack of significant mDA neuron death *in vitro* for many of the pesticides implicated by the PD PWAS. First, such pesticides may have no direct influence on PD, and the epidemiologic signal for association may simply have been driven by correlation with toxic co-exposures. Second, the pesticide influences PD through other disease-relevant pathways besides mDA toxicity, for instance, inducing pathological α-synuclein mis-folding with transsynaptic propagation[32], influences on the gut microbiome[33], or immune system disturbances[34]. Third, that key mechanisms were not recapitulated in our cell model. For instance, a metabolite was not generated during *in vitro* exposure to the purified mDA neurons that was actually the toxic species, as is the case for some organophosphorus pesticides that are activated to a toxic analog by cytochrome P450[35–37]. Finally, the pesticide could be toxic but only in combination with other pesticides.

We tested our platform’s capacity to reveal interesting synergistic exposures on the so-called “cotton cluster.” The cotton cluster is a group of highly correlated pesticides all associated with PD in the PWAS. Half of this particular cluster were defoliants applied almost exclusively on cotton. Analysis of co-treatment with components of this cotton cluster indicated that co-exposures involving trifluralin produced substantially more mDA cell death than any of the components alone (Figure 7D). For instance, the combination of trifluralin and tribufos produced more cell death than either alone and this pair caused more cell death than any other pairwise grouping. Taken together, the demonstration of individually mDA neuron-toxic pesticides and synergy with co-exposures suggests that eliminating overtly mDA toxic pesticide “hubs” from commonly used pesticide mixtures may reduce PD risk, even if not completely.

These findings with trifluralin prompted additional mechanistic work. Using functional assays in mDA neurons, we implicated mitochondrial mechanisms and mitochondrial dysfunction in trifluralin-mediated toxicity. Trifluralin is part of the dinitroaniline family of herbicides, known to cause disruption of cell division in plant cells and protozoa via de-polymerization of microtubules [38]. Trifluralin binds to tubulin in higher plants and protozoa (Leishmania [39], Trypanosoma[40] and Plasmodium [41]. Interestingly, no binding has been observed to mammalian or human tubulin [42]. Equally, we are not aware of any reports directly linking trifluralin to mitochondrial dysfunction in neurons, suggesting that the use of human dopamine neurons as a model can uncover novel toxicity pathways for pesticides. Links between trifluralin’s effect on microtubules and mitochondrial dysfunction, for example, are worthy of investigation. The results reported here strongly support an effect, whether direct or indirect, of trifluralin on neuronal respiration and mitochondrial function. These data are in keeping with an extensive literature documenting mitochondrial dysfunction is fundamental to PD [43–46]. Dysfunction in Complex I of the electron transport chain has been strongly associated with a PD phenotype in mouse models[47]. One caveat in our study is the relatively high dose of trifluralin used in our assays. Unquestionably, a challenge in the field is relating dose in a tractable timeframe in a cellular model with a chronic low exposure *in vivo*. In the future, more sensitive assays, for example imaging-based assays of mitochondrial potential or neuronal activity, will enable toxic effects of much lower doses of trifluralin to be assessed. As we continue research on co-exposures and mechanisms, we expect to further classify pertinent pesticides, including those our paradigm has already implicated, such as dicofol, the copper sulfates, and propargite.

Other potential limitations of our study are noteworthy. With regard to the exposure assessment for the epidemiologic study, we used a proximity model and record-based pesticide use reports from the study region (over 5.9 million reports in the study region). Without such a record system it would not be feasible to remotely assess 40 years of exposure to over 700 pesticides. Still, while our GIS-based ambient pesticide exposure assessment method is uniquely informative and has been previously validated [48], it allows for some level of exposure misclassification (i.e. error in the estimated exposure versus actual exposure), which is likely non-differential to case status and induces bias towards the null. This is described in more detail in the Supplemental Materials. Conversely, a great strength of this study is the record-based exposure assessment. The assessment did not rely on self-report, an advantage that cannot be overstated, especially in population-based research. One cannot rely on the general public in agricultural communities to know and report what pesticides are in their ambient environment, among thousands of applied products. Thus, this type of long-term, all-encompassing analysis in population-based PD research is dependent on a resource like the CA-PUR. Moreover, coupling the epidemiologic screen with experimental study, provides confidence in both the field and bench aspects of the paradigm.

Limitations of the *in vitro* modeling include: relative immaturity of cultured dopaminergic neurons; a lack of a blood-brain barrier (BBB); and a greatly simplified cellular environment lacking astrocytes, microglia, endothelial cells, and circulating factors. More advanced models, such as triculture systems that reconstruct the neuroimmune axis and “BBB on a chip” methods will permit investigation of more complex inter-cellular interactions and render the platform more physiologic [49, 50]. As noted above, dosing is an important consideration in our study. The pesticide doses used in our study, while comparable to other published reports, are high [4, 26, 27]. We used high doses to accelerate the onset of pathologies in the dish by using higher concentrations for shorter times rather than years worth of lower level exposure. Our *in vitro* models could also underestimate the toxicity of pesticides that require metabolism in the liver or by glial cells to generate harmful toxic metabolites. Additionally, with our live imaging system, we have chosen a straightforward and dramatic phenotype as the readout for pesticide toxicity--cell death. As noted above, more subtle phenotypes such as synuclein phosphorylation, abnormal neuronal activity, and dopamine oxidation will be important to detect effects of low doses of pesticides. With regard to the identification of trifluralin as a cause of mitochondrial dysregulation, more elaborated studies on the effect of nitroanilines (beyond trifluralin) in neuronal models of PD will be important. Little is known about the effect of other dinitroanilines herbicides developed as alternatives for trifluralin-resistant weeds. Such findings may have important implications for public health.

## CONCLUSIONS

The California Pesticide Use Report system has enabled us to begin to understand the breadth of pesticide application in agricultural regions and investigate how it relates to Parkinson’s disease in the community. Here, establishing a field-to-bench paradigm, we have combined record-based exposure assessment with a tractable and screenable iPSC-derived dopaminergic neuron model to identify and classify PD-relevant pesticides.

Our comprehensive, pesticide-wide association study has implicated both a variety of individual pesticides in PD risk and suggested relevant co-exposure profiles. Coupling this with direct testing *in vitro* on dopaminergic neurons, we have been able to pinpoint pesticides that were directly toxic to dopaminergic neurons. Further, real world co-exposure data has allowed us to develop co-exposure paradigms “in the dish” and establish which combinations of pesticides can indeed lead to greater, synergistic mDA toxicity. Ultimately, we have identified pesticides that are both ostensibly mDA-toxic and pesticides that are not mDA-toxic, but, nonetheless, associated with increased risk of PD. In time, our field-to-bench approach will enable us to further tie cellular pathologies back to this epidemiologic and environmental data, to mechanistically understand the individual and combinatorial effects of pesticides and their interactions with genetic susceptibility. Collectively, these studies will inform the judicious use of pesticides in agriculture.

## METHODS

### PEG Study Population

To assess pesticide and PD associations in the PWAS, we used the Parkinson’s Environment and Genes (PEG) study (n=829 PD patients; n=824 controls). PEG is a population-based PD casecontrol study conducted in three agricultural counties in Central California (Kern, Fresno, and Tulare)[17]. Participants were recruited and enrolled in two waves: wave 1 (PEG1): 2000-2007, n=357 PD patients, n=400 population-based controls; wave 2 (PEG2): 2009-2015, n=472 PD patients, n=424 population-based controls. Patients were early in their disease course at enrollment (mean PD duration at baseline 3.0 years (SD=2.6)), and all were seen by UCLA movement disorder specialists for inperson neurologic exams, many on multiple occasions, and confirmed as having idiopathic PD based on clinical characteristics[18]. Population-based controls for both study waves were required to be > 35 years of age, have lived within one of the three counties for at least 5 years before enrollment, and not have a diagnosis of PD. We identified potentially eligible controls initially through Medicare enrollee lists (2001) but mainly from publicly available residential tax-collector records (after 2001 due to HIPAA restrictions). More information about enrollment numbers can be found in the Supplemental Materials. Characteristics of the PEG study subjects are shown in Supplemental Table 12. The PD patients were on average slightly older than the controls and had a higher proportion of men, European ancestry, and never smokers.

### PEG Pesticide Exposure Assessment

We estimated ambient exposure to specific pesticide active ingredients (AIs) due to living or working near agricultural pesticide application, using record-based pesticide application data and a geographic information systems-based model[16]. We briefly describe our method, but provide more detail in the Supplemental Materials.

Since 1974, California law mandates the recording of all commercial agricultural pesticide use by pest control operators and all restricted pesticide use by anyone until 1989, and then (1990-current) all commercial agricultural pesticide use by anyone, to the PUR database of the California Department of Pesticide Regulation (CA-DPR). This database records the location of applications, which can be linked to the Public Land Survey System, poundage, type of crop, and acreage a pesticide has been applied on, as well as the method and date of application. We combined this database with maps of land-use and crop cover, providing a digital representation of historic land-use, to determine the pesticide applications at specific agricultural sites[51]. PEG participants provided lifetime residential and workplace address information, which we geocoded in a multi-step process[52]. For each pesticide in the PUR and each participant, we determined the pounds of pesticide applied per acre within a 500m buffer of each residential and workplace address each year since 1974, weighing the total poundage by the proportion of acreage treated (lbs/acre).

We were interested in long-term ambient exposures, and thus, considered the study exposure window as 1974 to 10 years prior to index date (PD diagnosis for cases or interview for controls) to account for a prodromal PD period. The exposure windows, which were on average 22 years for residential exposure and 18 years for workplace exposure, covered a very similar length and temporal period on average for patients and controls of each wave (study window comparisons shown in Supplemental Tables 1 and 12). To assess exposure across the study window of interest, for each pesticide, we averaged the annual lbs/acre estimates in the study window (e.g. lbs/acre estimates for all 22 years were averaged for participants with 22 years of exposure history), only using years for which the participants provided address information. This approach created one summary estimate of the average pounds of pesticide applied per acre per year within the 500m buffer for each pesticide, which was estimated at residential and workplace locations separately for each participant. We log transformed the estimates offset by one, centered, and scaled the estimates to their standard deviations (SD).

### Pesticide Regulation and Toxicity Classification

We linked each of the 288 pesticides included in the PWAS to chemical, regulatory, and toxicity information using publicly available databases. We used the Pesticide Action Network (PAN) database to determine the chemical class (e.g. organophosphorus) and use type (e.g. insecticide) for each pesticide[53]. We linked each pesticide associated with PD to registration status in the United States and European Union to highlight current, active agricultural use versus historical, past use. We obtained information on pesticide regulation using the U.S. EPA pesticide label database, U.S. EPA cancellation reports, California Product Label Database, and the European Commission database and Rotterdam Notifications. We further interrogated the PAN database to link information on known toxicity[53]. This includes whether the pesticide is drift prone based on vapor pressure, a groundwater contaminant, acutely toxic, a cholinesterase inhibitor, an endocrine disruptor, a carcinogen, or a developmental or reproductive toxicant. This information is based on public databases from government and international agencies, such as the EPA, California Prop 65 lists, World Health Organization, and International Agency for Research on Cancer. To identify a “most toxic” set of pesticides, PAN North America has designated certain pesticides as ‘bad actors’, information we also provide, if the pesticide meets any of the following criteria: known or probable carcinogen, reproductive or developmental toxicant, neurotoxic cholinesterase inhibitor, known groundwater contaminant, or a pesticide with high acute toxicity. Detailed methods on the toxicity and PAN designations can be found on the PAN website (https://www.pesticideinfo.org/).

### Statistical Analysis

A detailed description of analytic methods for the epidemiologic study, described in four parts, can be found in the Supplement, including methods used 1) to describe the extent of agricultural pesticide application in the study area; 2) for the pesticide-wide association analysis; 3) for pesticide group overrepresentation analysis; and 4) for PD-associated pesticide clustering. All analyses were done in R version 4.1.0.

### iPSC Reporter generation

Method for derivation of THtdtomato knockin clones was adapted from Ahfeldt et. al.[15]. Synuclein triplication iPSC lines derived from the patient used for this study have been described previously [13, 21]. Nucleofection of two plasmids containing sequence coding for gRNAs and a third targeting plasmid were co-nucleofected using the Amaxa 4D nucleofection system on a Lonza nucleofector. Targeting plasmid contained: the targeting vector with a TH homology arm followed by tdtomato, a 2A self-cleaving peptide sequence, a WPRE sequence, floxed puromycin selection cassette, and TH homology arm. Cells were allowed to recover for one day prior to initiation of puromycin selection. Surviving colonies were expanded and pooled for nucleofection of a pCAG-CRE:GFP vector allowing for Cre:LoxP based excision of the puromycin selection cassette and isolation by FACS of GFP+ cells. Cells were plated at clonal density and resulting colonies were expanded out and subcloned. A 5’ genotyping PCR producing a 626 bp product confirmed proper insertion of the 5 prime end of the reporter construct and a 3’ genotyping PCR producing an 878 bp product confirmed proper insertion of the 3 prime end of the reporter into exon 14 of the TH gene. PCR products were Sanger sequenced and aligned to the sequence expected after successful targeting. A third PCR designed to produce a 285 bp product was performed to evaluate the unmodified TH allele and assess for NHEJ errors. Colonies passing the above quality control were differentiated into dopaminergic neurons [8, 15, 54] and assessed for proper expression of THtdtomato insert with both live imaging and post-fixation immunoco-localization of tdtomato reporter (rabbit anti-RFP, Rockland, 1:500) with tyrosine hydroxylase (sheep anti-TH, Pel-Freeze #960101, 1:500). Neurons were fixed with 4% paraformaldehyde for 20 minutes at room temperature after 35 days of differentiation as embryoid bodies and 13 days in adherent culture. Clones that pass PCR, sequencing, and differentiation quality control steps are karyotyped to assess for any abnormalities.

### Dopaminergic Differentiation

Dopamine neurons were generated from the THtdtomato modified iPSCs in accordance with published protocols, using minimal modification[8, 15, 54]. iPSC cultures were maintained in growth media (StemFlex, mTESR plus, or mTESR media). On day 0 of differentiation, confluent iPSC cultures on matrigel or geltrex were dissociated into single cells using Accutase incubated for 5-7 minutes at 37 degrees following a 0.5mM EDTA in PBS wash. Accutase reaction was quenched with growth media and single cells were centrifuged, resuspended in growth media supplemented with Y-27632 at 10μM and FGF-2 at 20nM. Cells were plated at a density of 1×10^6 cells/mL in 15cm uncoated plates (30mL per plate) to form embryoid bodies. An additional 30mL of growth media is added the following day. On day 2, EBs are collected into 50mL conicals and spun at 200 x g for 3 minutes. Media is aspirated and EBs are replated in differentiation media (DMEM:F12 media mixed 1:1 with Neurobasal media, with PenStrep, B27 supplement without vitamin A, N2 supplement, 2-mercaptoethanol and glutamax) supplemented with 100nM LDN-193189 and 10μM SB431542. On day 3, 30mL of differentiation media is added with SAg (1μM) in addition to LDN-193189 and SB431542. Differentiation media is changed daily until day 5 at which point CHIR99021 is added (3μM) and media changes are then performed every other day. SB431542 and SAg are withdrawn on day 9. LDN-193189 is withdrawn on day 13 and BDNF (20ng/mL), GDNF (20ng/mL), TGFb3 (25ng/mL), dibutryl cAMP (0.5mM), and DAPT (10μM) are added. CHIR99021 is withdrawn on day 15 and TGFbeta3 is withdrawn on day 17. Spheres are maintained on uncoated plates with three times weekly media changes until dissociated for FACS and biochemical assays on day 35-42.

### Dissociation, FACS, and Live cell toxicant survival assay

Spheres are collected in a 15mL conical tube from suspension culture plate, washed with PBS and resuspended in 2mL of 0.25% Trypsin EDTA with 25ng/mL of DNASe added prior to incubation at 37 degrees in water bath or rotating shaker for 5-7 minutes. 500uL of FBS is then added to stop the reaction. Following a PBS wash, the EBs are tritutarted 5-7 times with a P1000 in a trituration solution (PBS with 5% FBS, 25mm Glucose, 1x glutamax). Cells are then washed with PBS and pelleted at 300 x g for 5 minutes 3-5 times prior to plating or FACS sorting. Large clumps and aggregates are filtered using a 35μM CellTrics filter. Y-27632 at 10μM is present for sorting and collection in differentiation media. A MoFlo Astrios and MoFlo XDP (Beckman-Coulter, both equipped with 100um nozzle at 30psi) were used to sort single THtdtomato + neurons based on scatter profile, pulse width, exclusion of Sytox Red dye, and tdtomato fluorescence. The brightest 30-40% of cells are included in order to minimize non-neuronal cell types or immature/neuronal progenitors expressing low level of the THtdtomato reporter. Sorted cells are plated onto polyornithine, poly-D-lysine, laminin, and fibronectin coated assay plates (Greiner 384 well plates) for survival assays. Sorted cells were plated at a density of 4×10^3 cells per well of a 384 well plate in a total volume of 45uL on the day of plating with Y-27632. Media wash and transition to Fluorobrite (GIBCO) live imaging media was performed the following day using an Apricot Personal Pipettor (Apricot Designs) leaving 90uL of fresh media per well. The first pesticide/toxicant treatment was performed two days after plating (Supplemental Figure 4). At five days after plating, half the media is aspirated, 45uL of fresh media are added and treatment is repeated. At nine days after plating, half the media as aspirated, 45uL of fresh media are added but treatment is not repeated. For additional quality control, live imaging is performed with a live cell chamber-equipped to maintain 5% CO2 and 37 degrees on an IXM High throughput microscope (Molecular Devices) using a 10x objective, a Texas Red filter (excitation 560/32nm; emission 624/40) and imaging four fields per well, resulting 72% coverage per well. Images were acquired immediately prior to treatment, at 7 days after first treatment and at 11 days after first treatment.

### Image analysis

Images were imported into Columbus (Perkin Elmer) analysis software. Differential brightness and roundness criteria permit the use of nuclei detection algorithms to detect the endogenous fluorescent reporter for accurate selection of cell soma with exclusion of neurites. The *Find Nuclei* script was used with method M, diameter was set at 13μM with splitting sensitivity of 0.15 and common threshold of 0.2. The nuclei detection algorithm was further refined to eliminate debris and doublets by selecting nuclei with area > 40μM and <400μM, and roundness >0.7. Additional brightness criteria (pixel intensity >500) were used to generate reproducible neuron cell body detection scripts that exclude debris and dead cells. Image analysis was performed in two steps to improve assay to assay reproducibility. The first step determined the average pixel intensity for detected cells that met size and roundness criteria. Average and standard deviation of fluorescence intensity was then determined for the control wells treated with DMSO. A second analysis was then designed to count all cells brighter than three standard deviations above the average fluorescence intensity in the control wells and measure neurites. These objects were counted as THtdtomato positive cells for analysis. Built in neurite detection algorithms were applied in order to detect neurites based on THtdtomato signal in these cellular processes using the *Find Neurites, CSIRO neurite analysis* method. For this neurite detection method, the following settings were used: smoothing window 3px, linear window 15px, contrast > 1, diameter >/= 7px, gap closure distance </= 5px, gap closure quality 0, debard length </= 15px, body thickening 5px and tree length </= 20px. Neurite analysis generated a sum of total neurite length per field analyzed which was used as the primary metric for neurite analysis.

### Toxicant Library generation

All compounds were ordered from Sigma Aldrich as the PESTANAL analytical standard whenever possible with the following exceptions: Tribufos (S,S,S-tributyl phosphorotrithioate) obtained from Fisher/Crescent Chemical; MSMA (Monosodium acid methane arsonate sesquihydrate) obtained from Santa Cruz biotechnology: oxydemeton-methyl obtained from Santa Cruz biotechnology. Based on available solubility data, compounds were dissolved in DMSO, water, or ethanol to a working dilution of 30mM. A subset had poor solubility that required a more dilute stock solution (15mM or less). A limited set of compounds identified in the PWAS analysis were omitted due to high dermal/inhalation toxicity in mammals, inability to obtain a highly pure formulation, or inadequate solubility in water, DMSO, or ethanol. Working stocks of compounds were pipetted into multiwell template plates and serial dilution was performed with an Apricot personal pipettor. Ethanol solubilized compounds were diluted 1:1 with DMSO to improve accuracy of pipetting and permit a parallel workflow to the DMSO and water plates. Individual compound plates were then generated from template plates following serial dilution and stored at −20 C until day of treatment. Apricot personal pipettor equipped with 125uL volume disposable tips was used to perform dilution of compounds in culture media and treatment of dopaminergic neurons.

### Combinatorial treatments

The combinatorial treatment plate maps were designed and performed using a D300e Digital Dispenser (Hewlett Packard) equipped with T8 and D4 dispenseheads. HP software was used to generate plate maps using the “synergy” function for 384 well format that produced all possible combinations of 6 different compounds at 10μM concentration. Compounds dissolved in ethanol were further diluted 1:1 with DMSO for dispensing accuracy. DMSO concentration was normalized to 0.3% DMSO to maintain accurate dispensing and account for higher order combinations of multiple compounds. DMSO controls were present on each plate to assess for extent of plate to plate variability within a given biological replicate. Upset plots were generated in R (Bioconductor) to visualize pesticide combinations on the x-axis (https://jokergoo.github.io/ComplexHeatmap-reference/book/upset-plot.html#upset-making-the-plot). Treatment timeline was identical to that described above for toxicant live imaging assays. Toxicants were combined as described. Synergy plots were created from an upset plot of each toxicant combination (ggupset(0.3.0)) and a bar plot of the day 11 neurons’ mean THtdtomato brightness (ggplot2 (3.3.5) with a viridis plasma palette (0.5.1). Each combination of two toxicant’s THtdtomato brightness values was compared via Student’s t-test, and p-values were adjusted for multiple testing with Benjaimin-Hochberg false discovery rate to q-values (reported in Supplemental Table 13).

### Agilent Seahorse XF Cell Mito stress assay

Cellular respiration of SNCA triplication THtd differentiated neurons in presence of ziram (45708 Milipore Sigma, CAS No 137-30-4) and trifluralin (111020171, MiliporeSigma, CAS No 1582-09-8) was accessed using the Seahorse XF 96 Extracellular Flux Analyzer and the XF Cell Mito stress test kit (Agilent Technologies) following Agilent Technologies guidelines. Mixed dissociated THtd neuronal cultures at day 35 of differentiation were seeded at a volume of 100μL and a density of 1×10^5 cells per well in the inner 60 wells of a PEI-laminin pre-coated Seahorse 96 well plate. The day of the experiment, 20 days after seeding (55 days of total differentiation), cells were treated with 0.3 % DMSO (control wells) and differentiation medium containing trifluralin at 90μM, 60μM and 30μM (treatment wells) for 6 hours. After this treatment period, medium was removed and 100μL the seahorse assay medium was added to each well of the entire plate to start the Mito stress assay, where Oligomycin (1 μM), FCCP (0.5 μM) and Rotenone/Antimycin-A (1 μM) are added at specific time points of the assay to measure metabolic outcome for each condition. After the assay finished, the assay medium was removed from the wells and cells were flash-frozen for subsequent measurement of total protein from each well using Pierce BCA assay kit (23225, Thermo Fisher). Total protein content was used to normalize the data. Data analysis performed with Wave software (Agilent Technologies).

### Mitochondrial function assays

The mitochondria subunit assay is based on the methodology published by Monzio et al. (Monzio et al. Stem Cell Reports 2018). *Cell pellet generation*: SNCA triplication differentiated neurons were dissociated from EBs at day 35 of differentiation. Mixed dissociated neuronal cultures were seeded at a density of 2×106 cells per well in the inner 8 wells of 24 well plates coated with polyethylenimine-Laminin (PEI-Lam). Cells were treated with trifluralin 30 days after seeding (65 days of total differentiation). Differentiation medium containing trifluralin at 90μM, 60μM 30μM, and 10μM and match 0.3% DMSO control were prepared. The old differentiation medium for each well to be treated was replaced with 1 ml of freshly prepared trifluralin containing differentiation medium. Cells under those conditions were incubated for 6 hours. After the incubation period, treated cells were harvested by detaching them from the wells by pipetting, transferred to 1.5mL Eppendorf tubes and centrifuged them at 4 °C for 5 min at 5000 rpm, medium was removed, and cell pellets were stored at −80 °C for biochemical analysis. *Protein Extraction*: Aqueous extraction of proteins from cell pellets was done by re-suspending the pellets in 100μL of 1X dilution from 4X blue LDS buffer (B0007, life technologies) plus protease (11697498001, Sigma) and phosphatase (4906837001, Sigma) inhibitors cocktail, sonication for 2x using a tip sonicator at 40 % power for 15 seconds on ice. Samples were boiled for 5 min at 100 °C and then centrifuged for 10 min at 850xg at 4 °C. Proteins contained in supernatant were quantified using the Pierce BCA assay kit (23225, Thermos Fisher). *Western blot*: Protein samples (30μg) were prepared using 4x Bolt LDS sample buffer (B0007, Thermos Fisher) and 10X Bolt Sample reducing agent (B0009, Thermos Fisher) and boil for 5 min and then loaded into precast Bolt 12-well mini Bis-Tris 4-12% gels and run in MES/SDS buffer for 30 min at 200 V. iBlot dry blotting system (Thermos Fisher) was used to transfer into iBlot nitrocellulose membrane (Thermos Fisher). Licor Odyssey Buffer PBS (Cat. no. 927-40000, Licor) was used to block the membranes at room temperature for 1 hr, followed by overnight incubation with agitation at 4 °C with the following primary antibodies: mitoProfile total OXPHOS Human WB antibody cocktail (1:1000, ab110411), SDHA monoclonal antibody (1:10000, 459200), anti-TOMM20 (1:1000, HPA011562) and actin antibody (1:1200, Sigma A2066). Four 5 min washes with PBS-T (0.005% tween) were performed followed by incubation for 1 hr at room temperature with secondary antibodies 680-anti-rabbit, 800-anti-mouse at 1:10,000 dilution in Licor Odyssey plus 0.1% Tween. Membranes were scanned using Licor Clx scanner. Analysis of protein bands was performed using image studio software.

## Contributions

*PUR, PWAS, co-exposure analysis*: Kim Paul

*Synuclein knock-in line generation, dopamine neuron experiments*: Richard Krolewski

*Mitochondrial assays, Seahorse assays*: Edinson Lucumi Moreno

*Transgenic reporter constructs*: Tim Ahfeldt

*Dopamine neuron experiments*: Jack Blank

*Statistics, graphing of Cotton cluster upset plots*: Kris Holton

*PWAS analysis interpretation/PUR map*: Melissa Furlong, Yu Yu

*GRAPES Exposure Assessment*: Myles Cockburn, Laura K Thompson,

*Neuro Exams/Data collection*: Jeff Bronstein

*Experimental design, data analysis, manuscript writing*: KP, RK, ELM, LR, VK, BR

## SUPPLEMENTARY MATERIALS

### PEG Study Enrollment

For PEG1 patient enrollment, 1,167 potentially eligible patients were identified through large medical groups, neurologists, and public service announcements, 604 did not meet eligibility criteria for the following reasons: 397 had been diagnosed with PD >3 years prior to recruitment, 134 lived outside the tri-counties, and 73 did not have PD. From the 563 remaining potential cases, 90 could not be examined by our movement disorder specialists, 56 declined or moved away, and 34 became too ill or died before the scheduled appointment. Of the 473 examined, 94 did not meet criteria for idiopathic PD, an additional 13 were reclassified as not having PD during follow-up, and 6 participants withdrew after examination and before interview. Of the remaining 360 patients, 357 provided all information necessary for inclusion in this study.

For PEG2 patient enrollment, we were able to draw study participants from the pilot PD registry program in California. This registry builds on a CA law enacted in 2004 to collect and register PD case records from all health care providers. Between 2010-2015, we screened 2,713 potentially eligible PD patients with an address in the tri-county study area reported to the registry. Of these, 397 denied having PD, 212 were diagnosed with PD prior to 2007 (the earliest diagnosis year allowed for enrollment), 39 did not live in the tri-county area, 1,042 were already deceased or too ill, and 293 were unable to be contacted or refused. Of the 730 eligible registry-reported patients, 601 were examined by the UCLA movement disorder specialists, and 126 did not meet criteria for idiopathic PD. Of the remaining 481 patients, 472 provided all information necessary for inclusion in this study.

Population-based controls for both study waves were required to be > 35 years of age, have lived within one of the three counties for at least 5 years before enrollment, and not have a diagnosis of PD. We identified potentially eligible controls initially through Medicare enrollee lists (2001) but mainly from publicly available residential tax-collector records (after 2001 due to HIPAA restrictions). We used two sampling strategies to increase enrollment success and representativeness of the source population: a) for PEG1, random selection from the Medicare enrollee lists and of residential parcels (identified from the tax-collector records) followed by mail or phone enrollment, and b) for PEG2, random selection of clustered households (five per cluster, identified through the tax-collector records) visited in person to enroll at least one eligible control from each cluster (only one per household allowed).

For PEG1, we contacted 1,212 potentially eligible controls. Of these individuals, 457 were ineligible: 409 were < 35 years of age, 44 were too ill to participate, and 4 resided primarily outside the study area. Of the 755 eligible population controls, 409 declined participation, were too ill, or moved before an interview was possible, resulting in the enrollment of 346 population controls. A pilot test mailing, for which the number of eligible participants who declined was not known, enrolled another 62 controls. Of the 408 PEG1 controls, 400 provided all information necessary for inclusion in this study. For PEG2, with the second sampling strategy, 4,756 individuals were screened, of whom 3,515 were ineligible (88% of these were out of the required age range) and 634 declined participation. Of the 607 PEG2 population controls enrolled, 183 completed only an abbreviated interview that assessed the most recent residential and workplace addresses, limiting long-term pesticide exposure assessment, thus, these individuals were excluded. From PEG2, 424 controls provided all information necessary for inclusion in this study.

### PEG Pesticide Exposure Assessment

We estimated ambient exposure to specific pesticide active ingredients (AIs) due to living or working near agricultural pesticide application, using record-based pesticide application data and a geographic information systems (GIS)-based model[16].

Since 1974, California law mandates the recording of all commercial agricultural pesticide use by pest control operators and all restricted pesticide use by anyone until 1989, and then (1990-current) all commercial agricultural pesticide use by anyone to the PUR database of the CA-DPR. This database records the location of applications, which can be linked to the Public Land Survey System (PLSS), poundage, type of crop, and acreage a pesticide has been applied on, as well as the method and date of application. The PUR database includes ~5.9 million records for the tri-county area and study period (1974-2017), documenting over 40 years of agricultural pesticide applications. We combined this database with maps of land-use and crop cover, providing a digital representation of historic land-use, to determine the pesticide applications at specific agricultural sites[51]. PEG participants provided lifetime residential and workplace address information, which we geocoded in a multi-step process[52]. For each pesticide in the PUR and each participant, we determined the pounds of pesticide applied per acre within a 500m buffer of each residential and workplace address each year since 1974, weighing the total poundage by the proportion of acreage treated (lbs/acre).

We were interested in long-term ambient exposures, and thus, considered the study exposure window as 1974 to 10 years prior to index date (PD diagnosis for cases or interview for controls), to account for a prodromal PD period. The exposure windows covered a very similar length and temporal period on average for patients and controls of each wave. For PEG1, the mean index year for PD patients was 2001.8 (SD=2.5 years) and 2002.4 (SD=1.6 years) for controls. For PEG2, the mean index year was 2009.2 (SD=3.3 years) for patients and 2009.2 (SD=0.5 years) for controls. This represents on average 22 years of ambient residential exposure information and 18 years of ambient workplace exposure information for study participants, taking the 10-year lag period prior to diagnosis or index date into account and only including years for which the participant reported an address we could geocode (comparisons shown in Supplemental Table 12 and the study windows are detailed in Supplemental Table 1). To assess exposure across the study window of interest, for each pesticide, we averaged the annual lbs/acre estimates in the study window (e.g. lbs/acre estimates for all 22 years were averaged for participants with 22 years of exposure history). For averaging across the study window, we only used years for which the participants provided address information.

This approach created one summary estimate of the average pounds of pesticide applied per acre per year within the 500m buffer for each pesticide. In the following, we refer to this measure as the intensity-weighted average (IWA). The IWA was estimated at residential and workplace locations separately for each participant. In total 722 pesticides had reported application within 500m of at least one PEG participant’s residence or workplace. However, we included only 288 chemicals in our PWAS, according to our criterion of having at least 25 study participants with estimated exposure in both the residence and workplace datasets. We log transformed the IWA, offset by one, centered and scaled the estimates to their standard deviations (SD).

### PEG Exposure Assessment LImitations

The exposure assessment method did not account for potentially relevant factors, such as wind patterns at the time of application or geographic features that may influence pesticide drift, and it also assumes that the participant was at the recorded location during the exposure relevant time or that the agent was still active and exposed the residents after application had occurred. However, being within a certain buffer of a pesticide application is one of the strongest predictors of air concentrations of pesticides, and resuspension from these applications is generally constant over at least a week-long period[55]. We have also previously validated our approach with high specificity for organochlorines with serum measurements[48].

### Statistical Analysis

We describe our epidemiologic analytic methods and results in four parts: 1) describing the extent of agricultural pesticide application in the study area; 2) the pesticide-wide association analysis; 3) pesticide group overrepresentation analysis; 4) PD-associated pesticide clustering. All analyses were done in R version 4.1.0.

First, we assessed the extent of agricultural pesticide application with basic descriptive statistics (mean, median, range, and standard deviation). We aggregated the PUR-reported total number of different pesticide active ingredients (AI) applied in both the entire study region (tri-counties: Kern, Fresno, and Tulare) and specifically within 500m of PEG participants’ residence and workplace locations, and we also aggregated the total reported pounds of pesticide applied. Pesticide products are often composed of one or more AIs, plus any inert ingredients which are generally proprietary and confidential. Thus, the 5.9 million PUR records in the tri-county area do contain information on different pesticide active ingredients applied at the same time or in the same product. However, the summed pounds of pesticide applied was specific to the AI within the product, and thus the way we aggregated did not over-count by summing the pounds of product applied.

Second, for the pesticide-wide association analysis, we conducted univariate, unconditional logistic regression to calculate odds ratios (ORs) and 95% confidence intervals (CIs) for PD with each pesticide (n=288) individually. Proximity estimated exposures at each location (residence and workplace) were assessed independently and separately for the PEG1 and PEG2 study waves. We combined the OR estimates from the study wave and location stratified analyses in a fixed effects meta-analysis, using a generic inverse-variance method for pooling (R, meta package, metagen function, coding as described Chapter 4.3[56]). We controlled for age, sex, race/ethnicity, education (years of schooling), and index year (year of diagnosis or interview) to account for temporal trends in pesticide use. We used a false discovery rate (FDR) correction to account for multiple testing. We also assessed differences in neighborhood SES based on census information, to account for confounding by SES factors that also vary with location. However, there were no differences in the neighborhood SES factors comparing where the PD patients versus the controls lived or worked. Thus, these factors were not included as covariates to control for confounding.

Third, we used overrepresentation analysis (ORA) to test for overrepresentation of pesticide groups (toxicity groups, chemical classes, and use types) in the set of PD-associated pesticides relative to all pesticides we assessed. Only 286 of the 288 tested pesticides are included in the ORA, as two pesticides could not be classified into groups. Specifically, ORA tests whether the “set of PD-associated pesticides”, meaning all pesticides associated with PD (considered at both FDR<0.05 and p<0.05), contains disproportionally more pesticides from a given group than expected given the distribution of the group in all pesticides assessed, tested with Fisher’s exact (R coding described Chapter 2.1[57]). This type of analysis is commonly applied to evaluate gene set overrepresentation. For the ORA, we considered all toxicity groups, classes, and use types that contained at least 3 pesticides from the PD-associated pesticide set. Altogether we assessed 8 toxicity groups, 5 chemical class groups and 9 use type groups, using the FDR<0.05 cut-off for the PD-associated pesticides, and 8 toxicity groups, 9 chemical classes, and 10 use type groups, p<0.05 cut-off, which are listed in the result tables (Supplemental Table 6).

Fourth, we assessed clustering of the PD-associated pesticides identified by logistic regression (FDR<0.10). We performed hierarchical Pearson correlation clustering analysis, separately for exposures at residential and workplace addresses, and provided a dendrogram with 1-R (Pearson) as the distance. As pesticides are regularly applied to the same field within the same season, year after year, this analysis assessing exposure clustering for PD-associated pesticides helps highlight real-world co-applications or mixtures and generates co-exposure profiles.

**Supplemental Figure 1.**
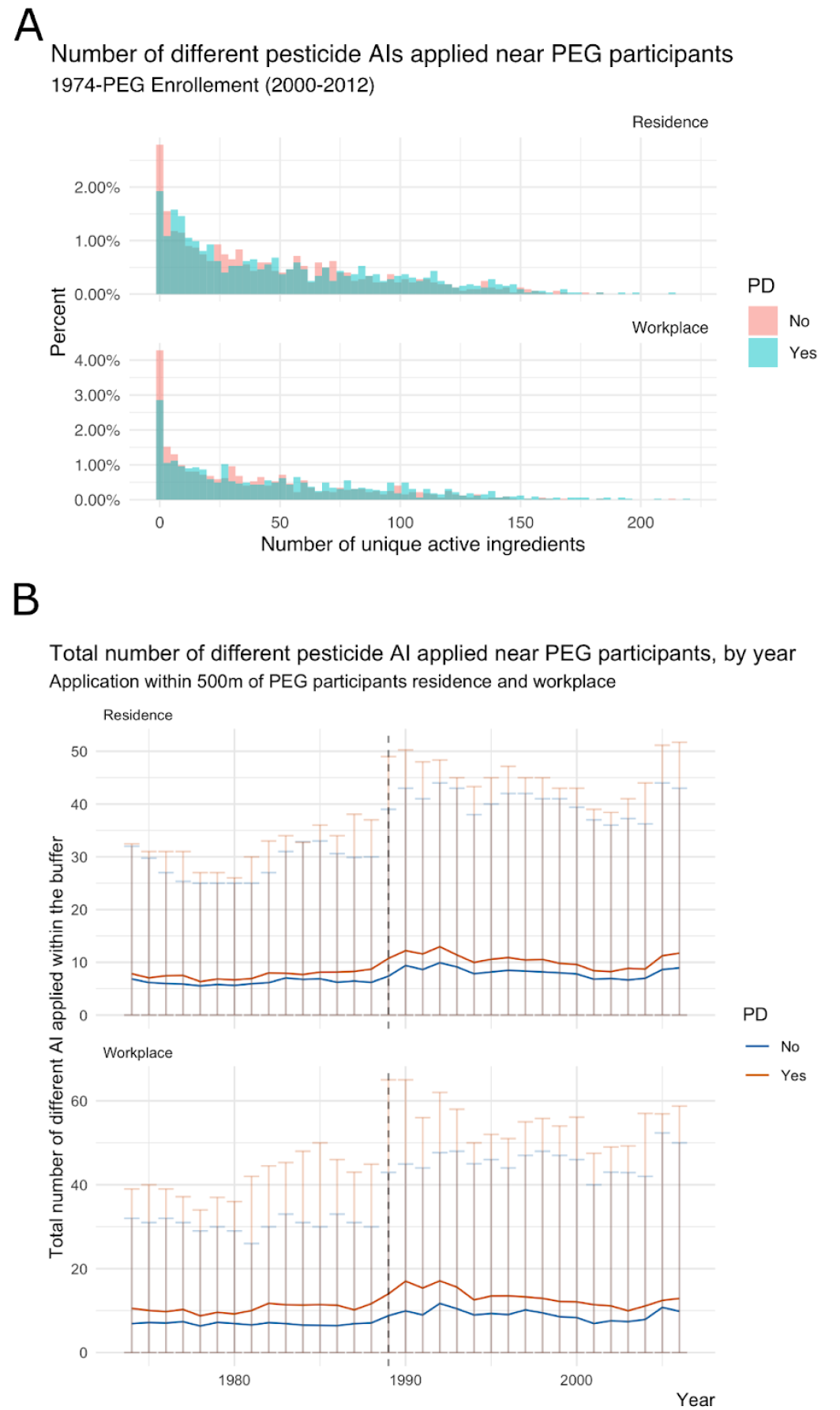
Total number of pesticide active ingredients applied near the PEG participants residence and workplace addresses (A) across the study window and (B) by year.

**Supplemental Figure 2.**
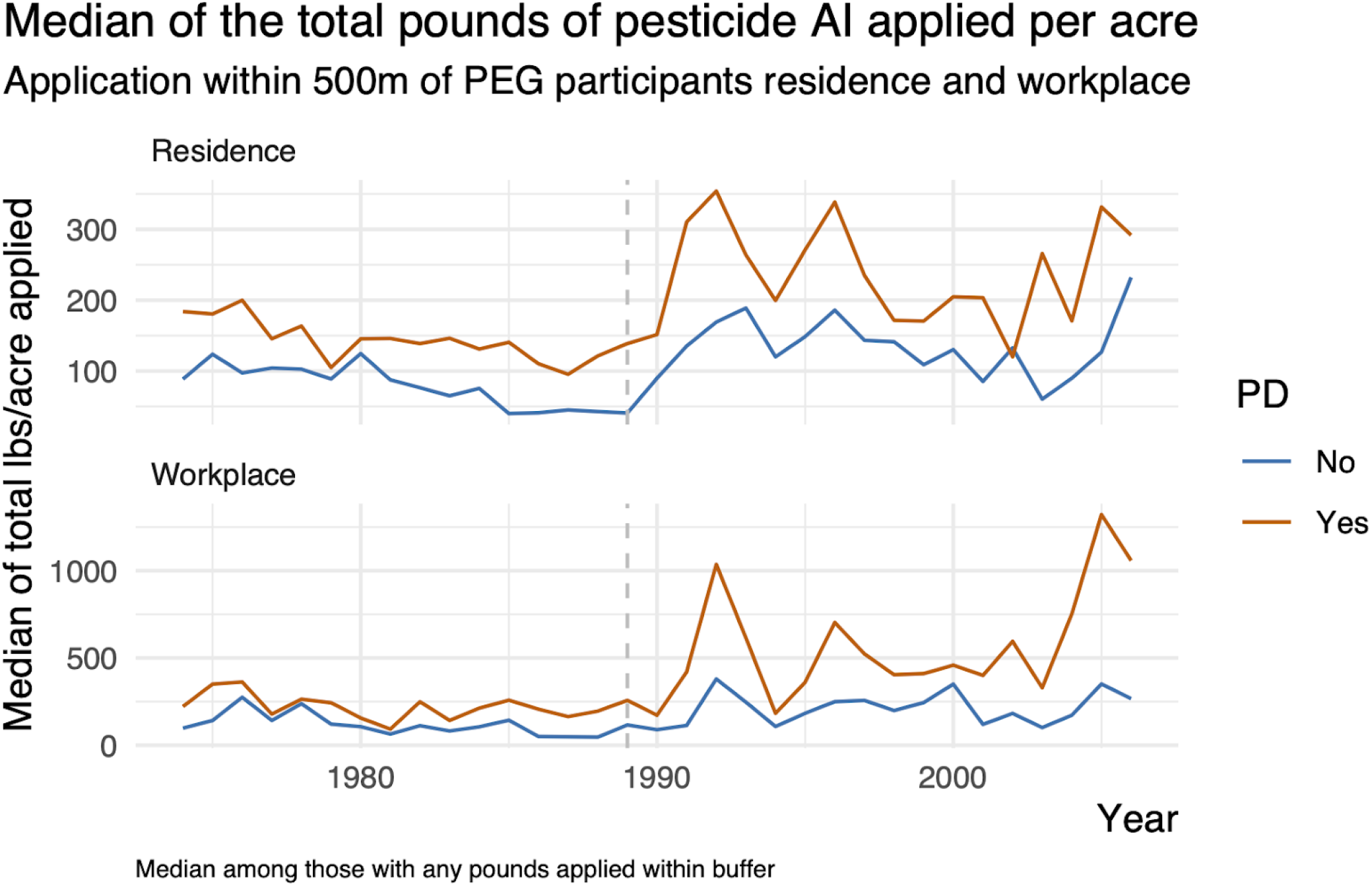
Median total pounds of pesticide active ingredient applied per acre near the PEG participants residence and workplace addresses.

**Supplementary Figure 3.**
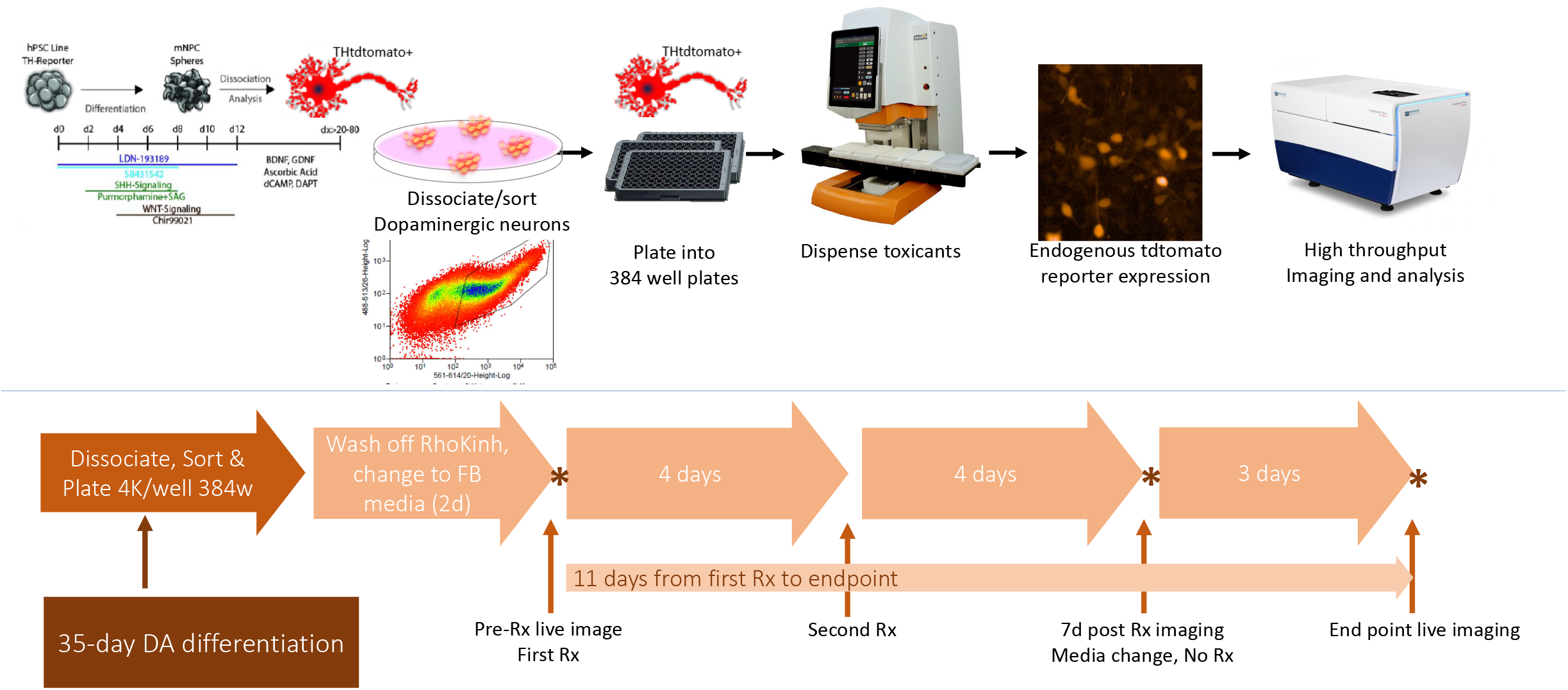
Dot plot displaying the odds ratio (OR) and 95% CI from the meta-analysis for all pesticides with an FDR<0.10 as well as results stratified by exposure location and PEG study wave.

**Supplementary Figure 4:**
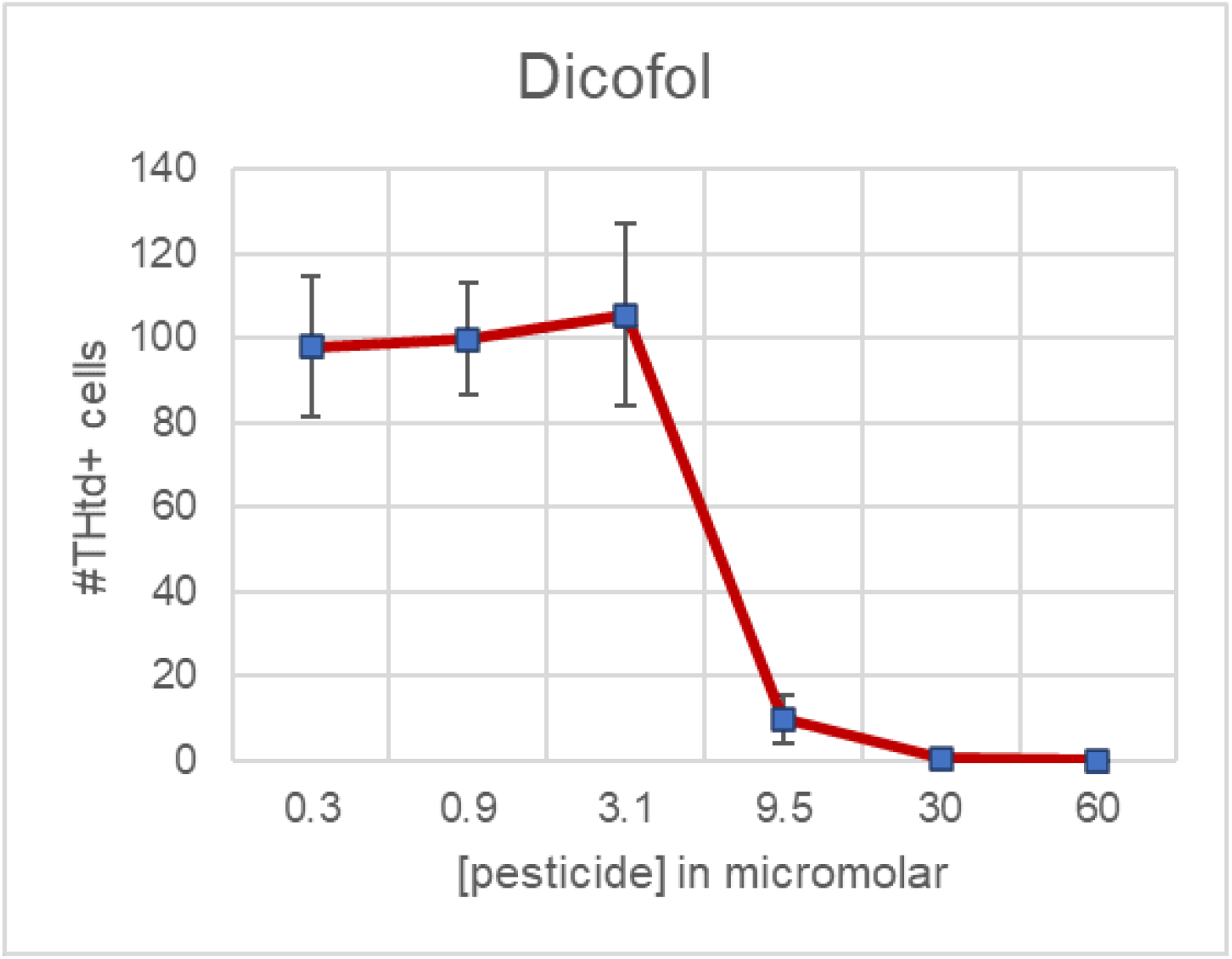
Direct treatment of mDA neurons and live imaging schematic. *Upper*. Assay design for custom PWAS in vitro dopaminergic neuron screen. Embryoid bodies/spheres were differentiated with a midbrain patterning protocol. Spheres containing THtdtomato+ dopamine neurons are dissociated for FACS between day 35-42 to purify THtdtomato+ cells which are then plated into multiwell assay plates using an Apricot Personal Pipettor, which was also utilized for media changes and toxicant treatment. Endogenous THtdtomato fluorescence was imaged on an IXM high content microscope fitted with a live cell chamber. *Lower*: timeline and sequencing of treatments, media changes, and imaging. Asterisks show imaging time points.

**Supplementary Figure 5:**
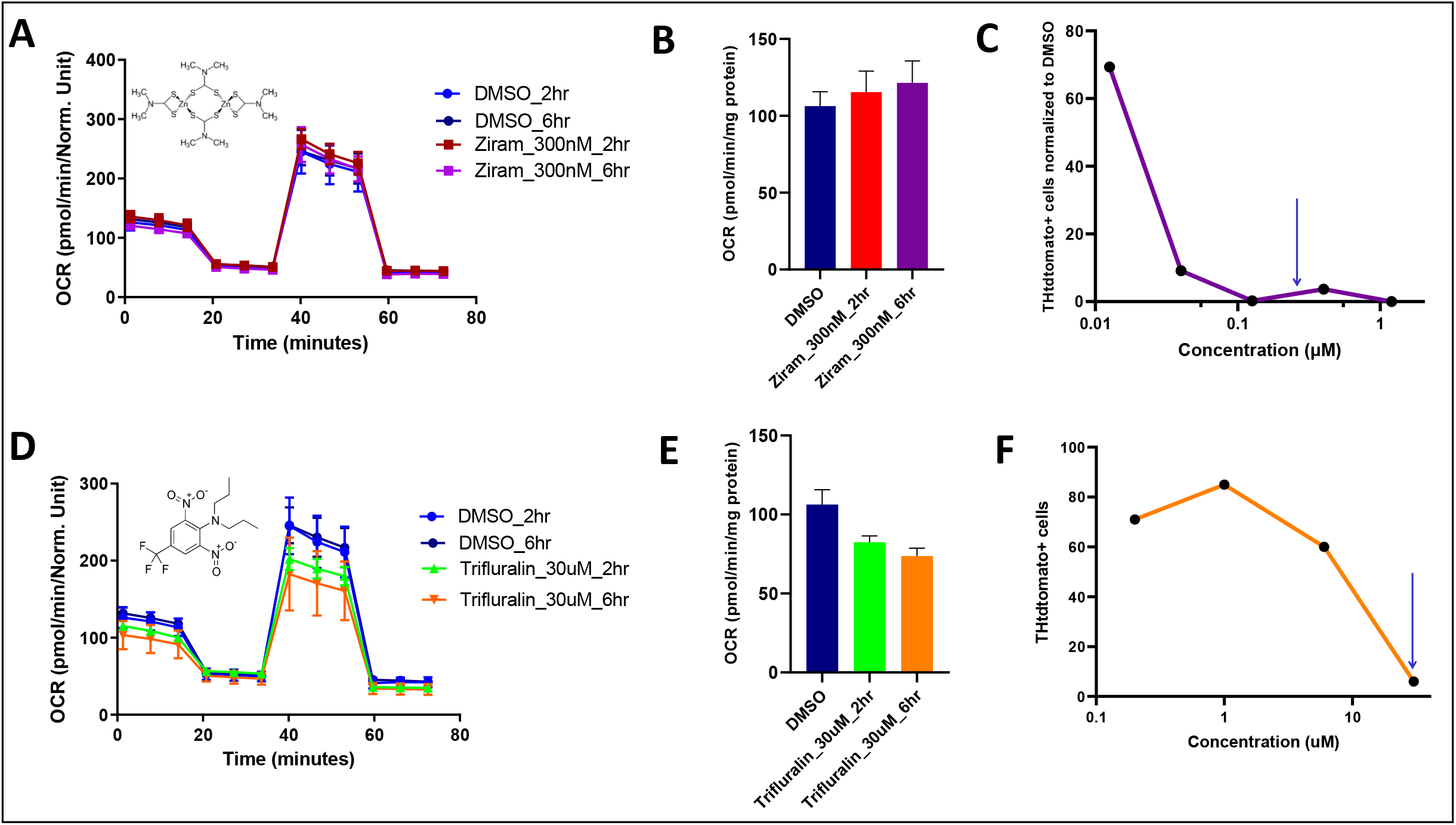
Dose response curve confirms toxicity of dicofol

**Supplementary Figure 6:** Comparison of Ziram and Trifluralin on oxygen consumption rates of SNCA-triplication differentiated neurons (**A**) Oxygen Consumption Rate curves on Agilent Seahorse XFCell Mito stress assay after treatment with 0.3 % DMSO and 300nM Ziram exposure for 2hr and 6 hr. (**B**) Spare respiratory capacity for the same conditions described in A. (**C**) Dose response curve for Ziram in a live cell survival assay similar to that described in Figure 5 demonstrating that Ziram dose used in Seahorse assay is adequate to kill dopamine neurons in the longer survival assay. (**D**) Oxygen Consumption Rate curves for 0.3 % DMSO and 30μM Trifluralin for 2hr and 6 hr. (**E**) Spare respiratory capacity for the conditions described in D. (**F**) Dose response curve for Trifluralin in a live cell survival assay similar to that described in Figure 5 demonstrating that Trifluralin dose used in Seahorse assay is adequate to kill dopamine neurons in the longer survival assay.

